# The transcription factor reservoir and chromatin landscape in activated plasmacytoid dendritic cells

**DOI:** 10.1101/2021.04.14.439791

**Authors:** Ritu Mann-Nüttel, Shafaqat Ali, Patrick Petzsch, Karl Köhrer, Judith Alferink, Stefanie Scheu

## Abstract

Transcription factors (TFs) control gene expression by direct binding to regulatory regions of target genes but also by impacting chromatin landscapes and thereby modulating DNA accessibility for other TFs. To date, the global TF reservoir in plasmacytoid dendritic cells (pDCs), a cell type with the unique capacity to produce unmatched amounts of type I interferons, has not been fully characterized. To fill this gap, we have performed a comprehensive analysis in naïve and TLR9-activated pDCs in a time course study covering early timepoints after stimulation (2h, 6h, 12h) integrating gene expression (RNA-Seq), chromatin landscape (ATAC-Seq) and Gene Ontology studies. We found that 70% of all described TFs are expressed in pDCs for at least one stimulation time point and that activation predominantly “turned on” the chromatin regions associated with TF genes. We hereby define the complete set of TLR9-regulated TFs in pDCs. Further, this study identifies the AP-1 family of TFs as potentially important but so far less well characterized regulators of pDC function.

## Introduction

Transcription factors (TFs) are known to bind to DNA-regulatory sequences to either enhance or inhibit gene transcription during cell differentiation, at steady state, and for exertion of cell effector functions (Vaquerizas et al., 2009; Wingender et al., 2018; Zhou et al., 2017). TFs also show unique expression patterns for different cell types and cellular states. The differentiation of distinct cell types from pluripotent stem cells is enabled by the expression of cell fate-determining TFs in progenitor cells. Transcription factors not only regulate cell development and effector functions by binding to *cis*-regulatory elements but also impact the accessibility of chromatin in different cell states (Serebreni and Stark, 2020). These latter TFs are called pioneering TFs and have the ability to remodel chromatin and thus modify the epigenome (Drouin, 2014). Chromatin is dynamically modified during cell differentiation leading to a cell-type specific landscape (Chauvistre and Sere, 2020; Deaton and Bird, 2011), which may be altered after cell activation. This process changes DNA accessibility for a particular set of TFs, that in turn modulate the expression of other genes important for cell identity and function. Efforts have been made to list and integrate all known mouse TFs in dedicated databases (db), such as Riken mouse TFdb (Kanamori et al., 2004) and TFCat (Fulton et al., 2009), amongst others. However, most of these were built before 2010 and have not been updated. The AnimalTFDB, most recently updated in 2019, classifies the mouse TF reservoir based on the structure of the DNA binding domains (Hu et al., 2019; Zhang et al., 2012). This database provides an accurate TF family assignment combined with TF binding site information in 22 animal species which also allows insight into TF evolution.

Plasmacytoid dendritic cells (pDCs) comprise a rare population of 0.2 to 0.8% of peripheral blood mononuclear cells (Liu, 2005). They were first described more than 40 years ago as natural interferon (IFN)-producing cells (IPCS) that activate NK cells after virus recognition (Trinchieri and Santoli, 1978). As we and others have shown, pDCs are now known for their capacity to produce unmatched amounts of type I IFN in response to stimulation of their toll like receptors (TLRs) (Ali et al., 2019; Asselin-Paturel et al., 2001; Bauer et al., 2016; Gilliet et al., 2008; Reizis, 2019). In contrast to other dendritic cell (DC) subsets, pDCs express only a limited repertoire of TLRs, namely predominantly TLR7 and TLR9 (Hornung et al., 2002), which recognize guanosine- and uridine-rich ssRNA and DNA containing CpG motifs (Diebold et al., 2004; Ishii and Akira, 2006; Wu et al., 2019). After TLR7 and TLR9 activation, in addition to type I IFN production, pDCs acquire the ability to prime T cell responses (Salio et al., 2004). CpG can be considered as an optimal and specific microbial stimulus for pDCs which induces TLR9 mediated signaling that leads to activation of IRF7 and NF-kB signaling pathways (Swiecki and Colonna, 2015). With regard to immunopathologies, unremitting production of type I IFN by pDCs has been reported in autoimmune diseases like systemic lupus erythematosus (Elkon and Wiedeman, 2012). Moreover, when recruited to the tumor microenvironment pDCs may induce immune tolerance and thus contribute to tumor progression (Le Mercier et al., 2013; Li et al., 2017). Thus, exploiting CpG for immunotherapeutic treatment to both enhance and repress pDC responses to mediate antitumor activity (Lou et al., 2011), treat allergy (Hayashi et al., 2004), and autoimmunity (Christensen et al., 2006) has been attempted in recent years. In addition, targeting specific TFs with the aim to control immunity and autoimmune disease (Lee et al., 2018) or to enhance cancer gene therapy (Libermann and Zerbini, 2006) has become the focus of attention in recent decades to develop immunomodulatory drugs.

Over the last years, different TFs have been determined as cell fate-instructive TFs in DCs. In particular, absence of the interferon regulatory factor 8 (IRF8) resulted in pDC-deficient mice (Tamura et al., 2005; Tsujimura et al., 2002). Bornstein *et al*. further identified IRF8 as an inducer of cell-specific chromatin changes in thousands of pDC enhancers (Bornstein et al., 2014). Further, mice deficient in the Ets family transcription factor Spi-B showed decreased pDC numbers in the bone marrow (BM) while pDC numbers were increased in the periphery. This indicated an involvement of Spi-B in pDC development, caused by a defective retainment of mature nondividing pDCs in the BM (Sasaki et al., 2012). In contrast to the phenotype of *Spi-B*-deficient mice, *Runx2*-deficient animals exhibited normal pDC development in the BM but reduced pDC numbers in the periphery due to a reduced egress of mature pDCs from the BM into the circulation (Chopin et al., 2016; Sawai et al., 2013). Finally, the *Tcf4*-encoded TF E2-2 is essentially required for pDC development as either its constitutive or inducible deletion in mice blocked pDC differentiation (Cisse et al., 2008). Using a combined approach to evaluate genome-wide expression and epigenetic marks a regulatory circuitry for pDC commitment within the overall DC subset specification has been devised (Lin et al., 2015). Even though the functions of selected cell fate TFs have been well described in pDCs, to our knowledge no global TF expression analysis after pDC activation has been performed for this cell type.

In the present study, we performed a detailed analysis on the changes in expression and chromatin accessibility for the complete set of all known TFs in pDCs in an early time course after activation. To this purpose, we used the AnimalTFDB data base and combined RNA-Seq, ATAC-Seq, and Gene Ontology analyses to define global TF gene expression, chromatin landscapes, and biological pathways in pDCs following activation. We defined epigenetic and transcriptional states using purified murine BM-derived Flt3-L cultured pDCs 2h, 6h, and 12h after TLR9 activation as compared to steady state. Based on our findings, we suggest a novel set of CpG-dependent TFs associated with pDC activation. We further identify the AP-1 family of TFs, which are so far less well characterized in pDC biology, as novel and possibly important players in these cells after activation.

## Results

### Expression of transcription factors in naïve and activated pDCs

To assess the impact of pDC activation on global TF expression in these cells, we simulated early events after virus infection in a time course study. To this end, we performed RNA-Seq of sorted BM-derived Flt3-L pDCs from C57BL/6N mice that were either left untreated or stimulated with CpG for 2h, 6h, or 12h. This synthetic double-strand DNA specifically activates endosomal TLR9 and is known to induce a robust type I IFN production (Gilliet et al., 2008). As the global definition of the mouse TF reservoir in this study we used 1,636 genes annotated by Hu *et al*. as TFs in the mouse genome (Hu et al., 2019). We evaluated the expression of all TFs in pDCs according to a formula by Chen *et al*., which takes into consideration the library length of the RNA-Seq run and the gene length to determine whether the gene is expressed or not (Chen et al., 2016). We found that 1,014 TFs (70% of all annotated TFs) are expressed in at least one condition, naïve or after TLR9 activation (2h, 6h, 12h) (**Fig. 1A**). The TFs expressed in pDCs were allocated to the different TF classes based on their DNA binding domain as described in the AnimalTFDB (Hu et al., 2019) (**Fig. 1B**). We found that more than half of all TFs (55%, 558 TFs in total) expressed in pDCs belong to the Zinc-coordinating TF group which use zinc ions to stabilize its folding and classically consist of two-stranded β-sheets and a short α-helix. Helix-turn-helix factors, of which 158 (16%) were expressed in pDCs under the defined conditions, comprise several helices mediating multiple functions such as insertion into a major DNA groove, stabilization of the backbone and binding to the overall structure of the DNA (Aravind et al., 2005). Furthermore, 10% (104 TFs) of all TFs expressed in pDCs belong to the Basic Domain group, which contains TFs that become α-helically folded upon DNA binding (Patel et al., 1990; Weiss et al., 1990). 44 expressed TFs (4%) belong to the Other α-Helix group exhibiting α-helically structured interfaces are required for DNA binding. In addition, 32 of the TFs (3%) found in pDCs are β-Scaffold factors which use a large β-sheet surface to recognize DNA by binding in the minor groove. Lastly, another ~100 TFs (12%) were of unclassified structure, meaning their mode of action for DNA binding is unknown. Strikingly, some TF families were not expressed in pDCs at all (**Fig. 1C**), such as the AP-2 family in the Basic Domain group, the GCM family in the β-Scaffold group, the Orthodenticle homeobox (Otx) TFs in the Helix-turn-helix group, Steroidgenic factor (SF)-like factors in the Zinc-coordinating group, and the DM group, first discovered in *Drosophila melanogaster*, among the unclassified TFs. Other TF families showed expression of all family members in at least one condition (steady state, or CpG 2h, 6h, 12h), such as the Transforming growth factor-β stimulated clone-22 (TSC22) family in the Basic Domain group, Runt and Signal Transducers and Activators of Transcription (STAT) factors from the β-scaffold classification, and E2F and Serum response factor (SRF) factors in the Helix-turn-helix group. In summary, 70% of all genes annotated as TFs in the mouse genome (1,014 out of 1,636) were expressed either in naïve or activated pDCs (CpG 2h, 6h, 12h), covering a wide range of TF classes based on different DNA binding mechanisms.

**Fig. 1.**
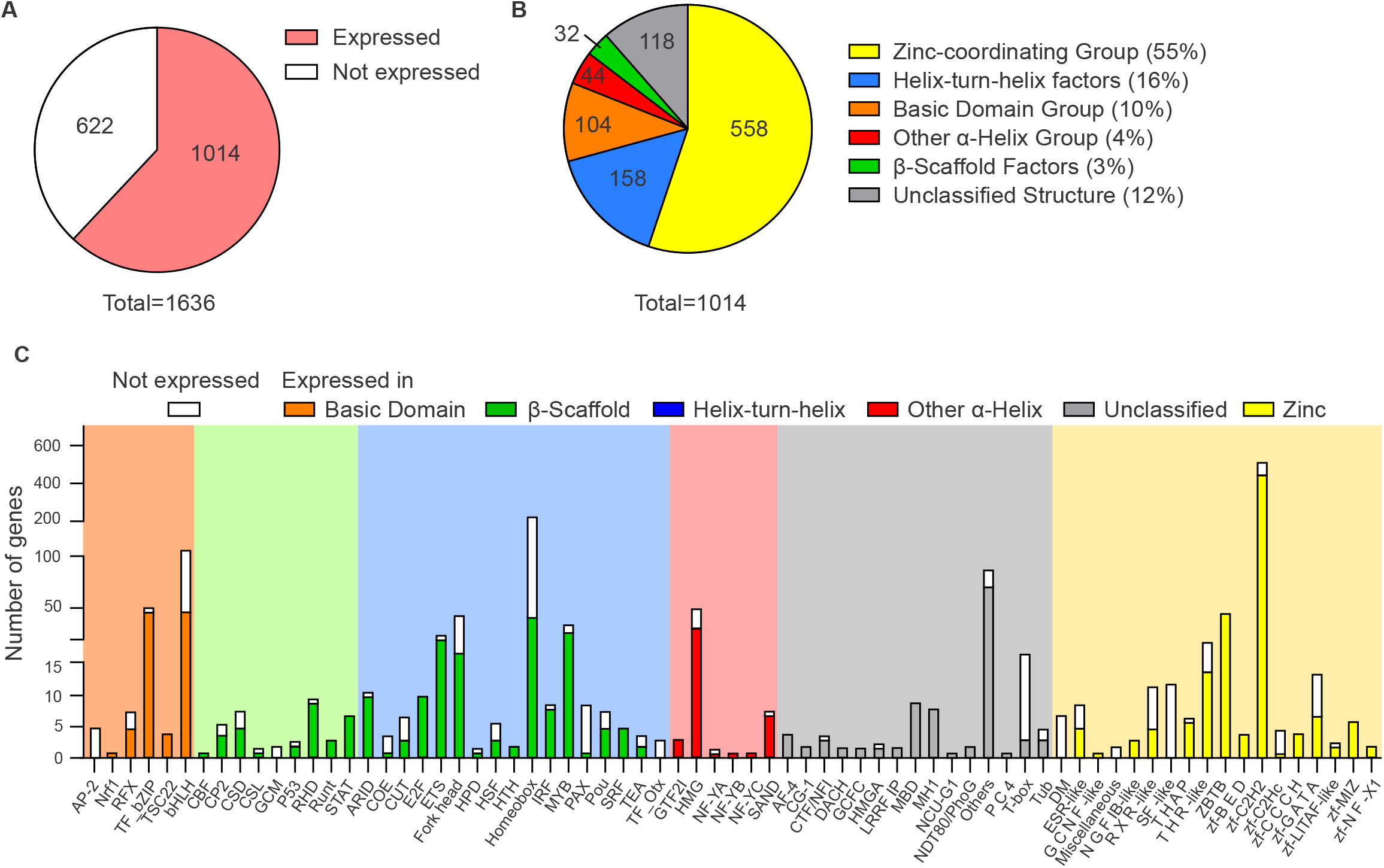
Expression of transcription factors in pDCs. **A** Expression of TFs in pDCs in at least one of the following conditions: naïve, CpG 2h, 6h or 12h (n=3 per condition). **B** Categorization of the expressed TFs according to Hu *et al*. (Hu et al., 2019). **C** Number of expressed vs non-expressed genes per TF family of a TF class is plotted.

### Activation-dependent TF expression changes

We next investigated the impact of pDC activation on changes in expression of TFs using our time course RNA-Seq study. The similarity of our biological replicates in each condition was evaluated with a Pearson correlation analysis. Our results revealed high similarity (<95%) for the biological replicates used in the respective conditions of the RNA-Seq data set. Notably, the differences in the Pearson correlation coefficient between the naïve and first stimulation time point (CpG 2h) were higher than the differences observed between the later CpG stimulation time points (6h, 12h) (**Fig. 2A**). We used the data for differential expression analysis of genes between pDC states, not only comparing TF expression levels between different CpG stimulation time points vs steady state but also between the different CpG stimulation time points between each other (**Fig. 2B**). The total number of differentially expressed TFs (DETFs) with a fold change |FC|>2 and a p<0.05 between stimulated vs naïve pDCs (452 DETFs in 2h vs 0h; 400 DETFs in 6h vs 0h; 335 DETFs in 12h vs 0h) was higher than the absolute number of TFs showing expression changes between the CpG conditions (270 DETFs in 6h vs 2h; 119 DETFs in 12h vs 6h; 358 DETFs in 12h vs 2h). This reflects the results from the Pearson correlation analysis (**Fig. 2A**). Interestingly, by comparing TF gene expression in 2h stimulated vs unstimulated pDCs, a higher number of TF genes were down-regulated in expression after TLR9 stimulation than were upregulated in these cells (271 vs 181). With increased duration of pDC stimulation, the difference in the number of TFs that were up-vs down-regulated diminished (208 down vs 192 up in 6h vs 0h). Finally, at the longest stimulation time used in this study (12h vs 0h), the number of up-regulated TF genes was higher than the number of down-regulated TF genes (179 vs 156). Comparing the CpG stimulated samples amongst each other, more TFs exhibited increased expression with longer stimulation times than there were TFs showing reduced expression levels (171 up vs 99 down in 6h vs 2h; 63 up vs 56 down in 12h vs 6h; 234 up vs 124 down in 12h vs 2h) (**Fig. 2B** and **C**). In total, we identified 661 unique TF genes that are differentially expressed between at least one of the compared pDC states |FC|>2, p<0.05, pDC at steady state, or after CpG activation at 2h, 6h, 12h). To evaluate patterns of expression changes for all 661 differentially expressed TFs, we next carried out hierarchical clustering of all TF genes based on the normalized expression in naïve and stimulated pDCs (**Fig. 2B**). This led to the definition of five different clusters of TFs according to their expression pattern (**Fig. 2D**). Cluster I, IV and V contained TFs with large expression changes after short duration of pDC stimulation (2h), while cluster II and III contained TFs that exhibit altered expression only with longer duration of cell stimulation (6h, 12h). Cluster V contained genes that were all down-regulated at any time point after CpG stimulation as compared to the unstimulated condition (**Fig. 2D**). In more detail, TFs driving either pDC (e.g. *Tcf4, Spib, Runx2*) or classical DC (cDC) (e.g. *Nfil3, Spi1, Id2*) development (Bornstein et al., 2014; Sasaki et al., 2012; Sawai et al., 2013; Tamura et al., 2005; Tsujimura et al., 2002) were distributed over all clusters I to V. This highlights variable expression patterns of DC cell fate TFs after pDC activation. In summary, in this time course study that models early events after virus infection, we identified in total 661 unique CpG-dependent TF genes that show significant differential expression in at least one condition compared to another |FC|>2, p<0.05, pDC at steady state, or after CpG activation at 2h, 6h, 12h). Further, pDC activation showed time dependent activating as well as inhibiting effects on the expression of TFs.

**Fig. 2.**
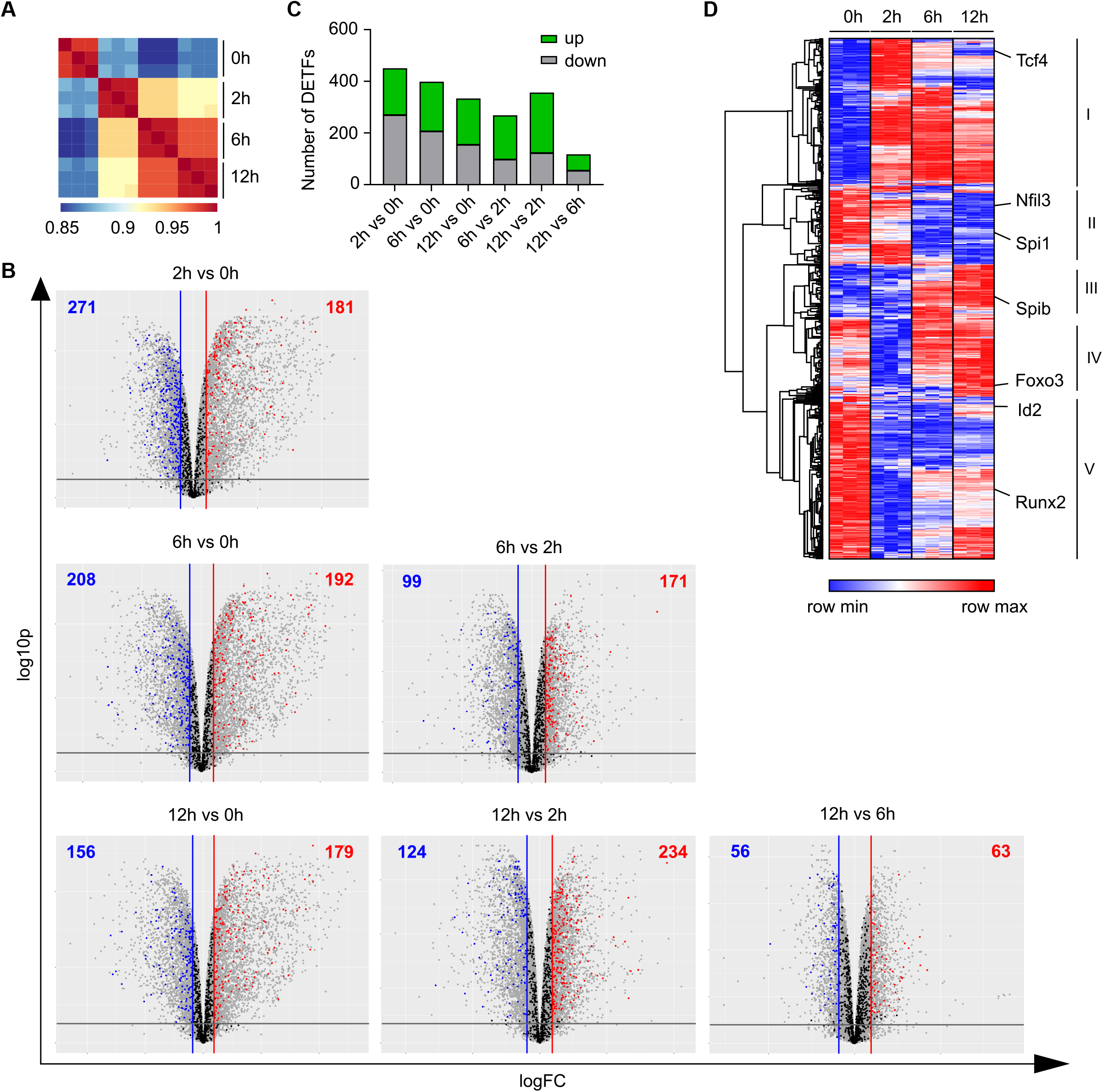
RNA-Seq reveals significant TF expression changes after pDC activation. **A** Pearson correlation plot for samples used in RNA-Seq. pDCs (CD3^-^CD19^-^CD11c^+^CD11b^low^B220^+^SiglecH^+^CD317^+^) were sorted from BM-derived Flt3-L cultures of C57BL/6N mice and cells were left either naïve or stimulated with CpG for 2h, 6h or 12h. **B** Volcano plots showing global expression of genes in sorted pDCs at steady state and after 2h, 6h, and 12h of CpG stimulation. TF genes with a |FC|>2 and a p-value of <0.05 corrected for the false discovery rate (FDR) were considered significantly differentially expressed and are marked in colour (red and blue). **C** Heatmap showing normalized expression values (cpm, count per million) of differentially expressed TF genes from (**B**) in pDCs at steady state and after 2h, 6h, and 12h of CpG stimulation. Hierarchical clustering on rows with average linkage and the One minus Pearson correlation metric was performed.

### Gene ontology analysis of CpG-dependent TFs

Next, downstream gene ontology (GO) analyses of RNA-Seq data were performed to unravel the biological processes in which CpG-dependent TFs are involved. For this purpose, functional annotation clustering with the 661 TF encoding genes defined as CpG-dependent |FC|>2, p<0.05) was performed on DAVID including terms for biological processes (BP), molecular functions (MF), and cellular components (CC). The analysis produced 16 clusters, out of which the 9 non-redundant and most relevant in the context of innate immunity are depicted in **Fig. 3A** (complete list in **Table S1**). The GO analyses produced an individual fold enrichment for each GO term (**Fig. 3A**, right column), and in addition, an enrichment score for each cluster containing several GO terms (**Table S1**). The order of the clusters from top to bottom follows a decrease in the cluster enrichment score, establishing a hierarchy of importance for the biological processes affected. Cluster one contained GO terms for DNA binding, transcription, and nuclear localization with a ~5 fold enrichment comprising more than 400 genes in each term. This confirmed the inherent DNA binding capacity of the defined murine TF reservoir by Hu et al. (Hu et al., 2019) and proved the applicability of our approach. The following clusters comprised less than 25 unique genes per GO term but significant fold enrichments for most GO terms drawing attention to specific TFs involved in particular biological processes in pDC activation. Cluster 2 contained GO terms associated with the circadian rhythm and regulation of gene expression (e.g. *Klf10, Jun*). We further found GO terms enriched for the IκB/NFκB complex, NIK/NFκB signaling, and IκB kinase/NFκB signaling (e.g. *Nfkb1, Nfkb2, Rel*), which showed the highest fold enrichment (up to 25 fold) among all GO terms and clusters. In line with this, it is well known that CpG activates the canonical TLR9-Myd88-NFκB/IRF7 signaling pathway in pDCs (Tomasello et al., 2018). Another cluster contained processes involving SMAD proteins (e.g. *Smad1, Smad2, Smad3*), signal transducers for TGFβ receptors, involved in receptor binding, signal transduction, and protein complex assembly. Of note, it is known that pDCs exposed to TGFβ lose their ability to produce type I IFN after TLR9 stimulation (Saas and Perruche, 2012). Another significantly enriched cluster comprised GO terms for various processes involving the endoplasmic reticulum (e.g. *Cebpb, Ddit3*), an important site of intracellular protein and lipid assembly. GO terms containing TFs that regulate sumoylation (e.g. *Pias4, Egr2*), posttranslational modifications that e.g. coordinate the repression of inflammatory gene expression during innate sensing (Decque et al., 2016), were also significantly enriched and clustered together. As expected, CpG-dependent TFs were enriched in GO terms for the JAK-STAT signaling pathway (e.g. *Stat1, Stat2, Stat3*) activated by binding of type I IFN to the type I IFN receptor. TFs affecting mRNA binding processes (e.g. *Mbd2, Ybx2*) which are required for synthesizing proteins at the ribosomes, were also affected. The fact that epigenetic modulators (e.g. *Prdm9, Kmt2c*) were enriched, highlights the importance of gene expression regulation of TFs in pDCs by modifications that alter the physical structure of the DNA after CpG stimulation. In summary, we find that CpG-dependent TFs are involved in a wide variety of biological processes, such as circadian regulation, mRNA binding, and signaling pathways such as the NFκB and JAK-STAT pathways. The analyses revealed the importance of these biological processes being affected by pDC activation in a hierarchical manner according to their attributed relevance. This opens up the opportunity to investigate specific TFs involved in processes that have not been fully elucidated for pDC biology.

**Fig. 3.**
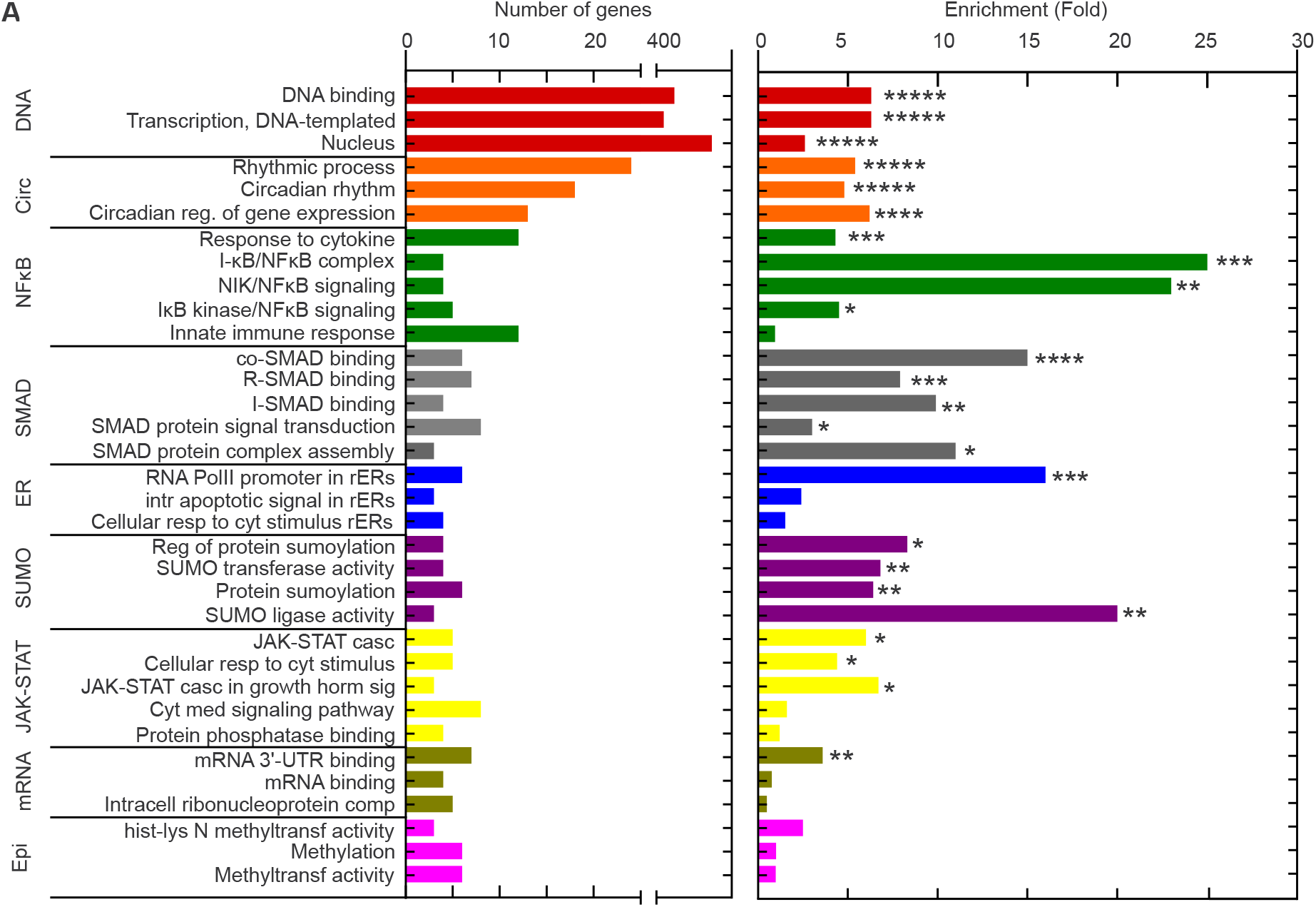
Gene Ontology analysis of CpG-dependent TFs. 661 CpG-dependent TFs (|FC|>2, p<0.05) were analysed by DAVID functional annotation to produce gene clusters (>2 genes/cluster) corresponding to biological process (BP), molecular function (MF), and cellular component (CC) GO annotation terms. Those significantly associated with the TF gene list are plotted with the numbers of genes for each term along with the fold enrichment for each term. A few terms were excluded as being redundant or having wider meaning (**Table S1**). Abbreviations are as follows: casc = cascade; cyt = cytokine; horm = hormone; med = mediated; reg = regulation; rERs = response to endoplasmic reticulum stress; resp = response; sig = signaling.

### pDC activation modulates chromatin accessibility for binding of TF families

Another hallmark of cell activation is the modification of the chromatin landscape. To better understand how the chromatin accessibility of different TF families is altered in pDCs in the course of activation, we performed ATAC-Seq in naïve and 2h CpG activated pDCs. Pearson correlation analysis for the ATAC-Seq data reveals >95% similarity for all biological replicates (**Fig. 4A**). A quantitative analysis of peak intensities across sample conditions and a differential analysis to determine the number and regions of activation-dependent accessible chromatin peaks was performed. Comparing the specific genomic locations such as introns, 3’-UTRs, distal (1-3kb) and proximal (0-1kb) promoter regions with accessible chromatin between naïve and 2h CpG stimulated pDCs, we found that chromatin is mostly open in distal intergenic and intron regions in both conditions. However, there was no apparent shift in the distribution of genomic locations where chromatin is accessible in pDCs after cell activation (**Fig. 4B**). This suggests that TLR9 activation regulates the chromatin accessibility globally in pDCs but does not induce shifts in the chromatin landscape per se. Overall, we detected ~116,000 accessible regions (peaks) across samples in naïve and activated states. Next, we performed a differential analysis using the DESeq2 algorithm to quantify the number of CpG-dependent accessible peaks. pDC activation substantially altered the chromatin landscape leading to ~16,600 altered accessible regions (|FC|>2, p<0.05, **Fig.4C, D**). In detail, 2h CpG stimulation of pDCs resulted in 13,226 peaks with increased accessibility and 3,381 peaks with decreased accessibility (**Fig. 4C, D**). Roughly 80% of all CpG-dependent chromatin regions in 2h stimulated pDCs exhibited increased DNA accessibility as compared to naïve pDCs. This suggests that more of the pDC chromatin landscape is turned on” rather than being “turned off” after pDC activation. To unravel the biological significance of the activation-dependent chromatin states for the more accessible vs the less accessible DNA regions in pDCs, a differential motif analysis using the HOCOMOCO database (Kulakovskiy et al., 2018) was performed (**Fig. 4E**). The purpose of the analysis was to identify TF families that gain or lose access to DNA after pDC activation which would hint at pathways being affected after activation. At the same time, this unbiased approach allows the identification of TFs that have not been associated with this cell type before. This motif analysis revealed that TFs belonging to the JAK-STAT and the NFκB signaling pathway have increased accessibility to their specific DNA binding regions after CpG stimulation. Besides the NFκB family, we identified the AP-1 family of TFs as one of the most significant hits to gain access to the DNA in our search. This type of TF remains so far less well characterized in pDCs after pathogen encounter or in pDC-specific functions in chronic inflammatory or autoimmune disorders. Albeit the AP-1 member c-Fos has been shown to be required for type I IFN induction, a hallmark function of pDCs, in osteoclast precursor cells after RANKL treatment (Takayanagi et al., 2002). On the other hand, Ets family members belonging to the Helix-turn-helix family of TFs and Zinc-coordinating zf-C2H2 TFs had less access to DNA. Strikingly, pDC-driving cell fate TFs such as IRF8 and RUNX2 showed motif enrichment in two sets of regions, one set with increased and another set with decreased chromatin accessibility after pDC activation. Hence, pDC-driving cell fate TFs both gained and lost access to specific DNA regions after TLR9 activation. We next performed a more detailed analysis searching for enrichment of TF motifs among all regions that contain the promoter sequence of one or more genes. As TFs can regulate gene expression by binding to the promoter site of genes this analysis hints at TF families that exert a functional binding occupancy in the investigated chromatin regions. We previously determined that 13,226 regions exhibit increased chromatin accessibility after pDC activation. Out of these, 2,174 regions were associated with the promoter of one or more genes. An unbiased motif enrichment search revealed that TFs belonging to the NFκB family (e.g. NFκB1, NFκB2, TF65), the AP-1 family (e.g. ATF3, JUN, FOSB), and the JAK-STAT family (e.g. STAT1, STAT2), as well as pDC cell fate TFs (e.g. RUNX2, IRF8) are among the top hits for TFs with DNA binding domains present in promoter associated chromatin regions which gain accessibility after pDC activation (**Table S2**). In summary, the differences in chromatin landscapes of naïve and 2h CpG stimulated pDCs point to a substantial amount of epigenetic modulation of thousands of pDC regions. Also, these analyses unravelled the AP-1 family of TFs, which have so far been less well characterized in pDC biology, as possibly important players in these cells after activation.

**Fig. 4.**
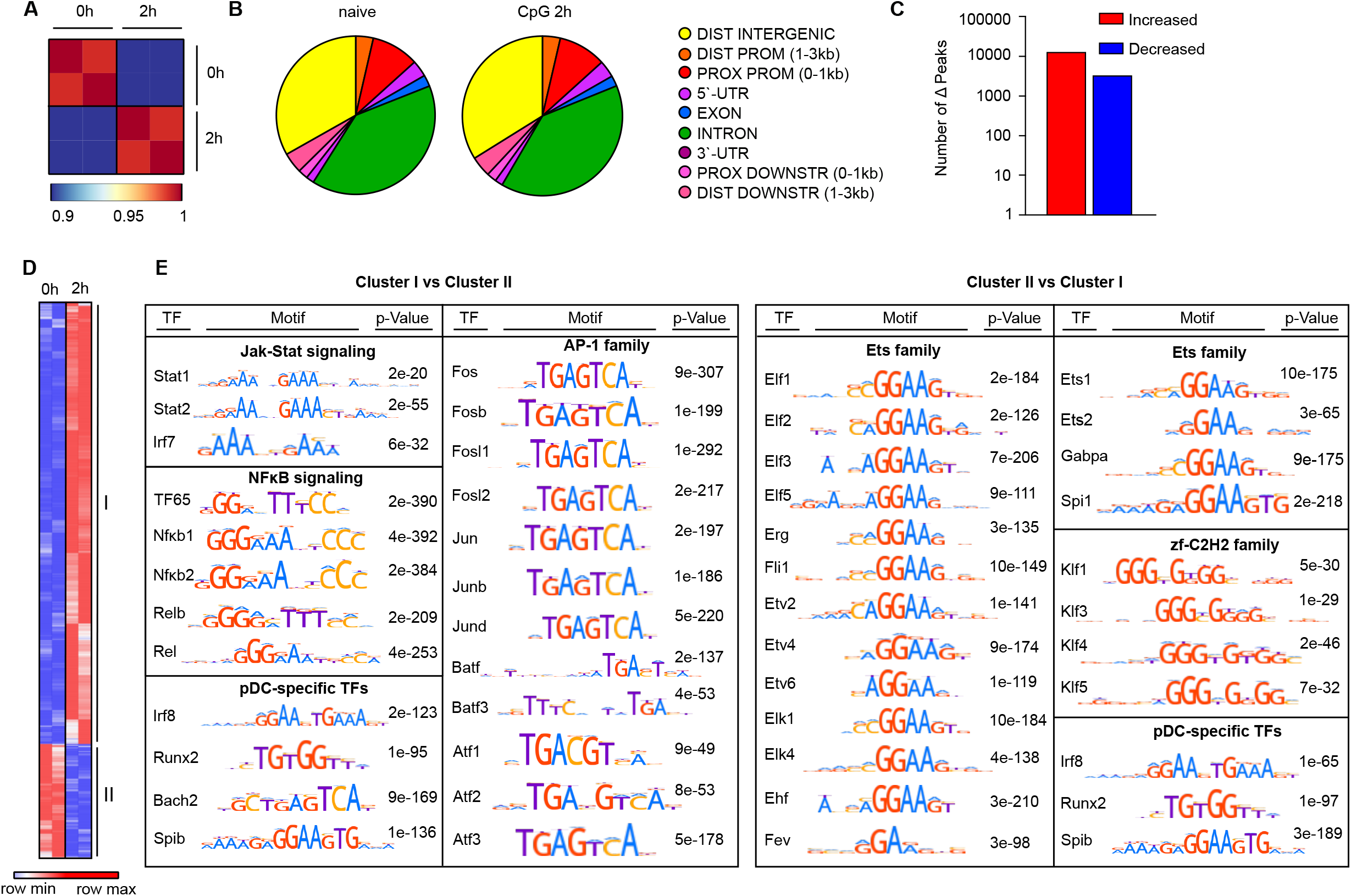
pDC activation increases and decreases chromatin accessibility of thousands of regions. **A** Pearson correlation plot for samples used in ATAC-Seq. pDCs (CD3^-^CD19^-^CD11c^+^CD11b^low^B220^+^SiglecH^+^CD317^+^) were sorted from BM-derived Flt3-L cultures of C57BL/6N mice and cells were left either naïve or stimulated with CpG for 2h (n=2). **B** Genomic location distribution of open chromatin sites in naïve and CpG stimulated pDCs according to ATAC-Seq. Two biological replicates were used per condition, and results are shown for pooled samples per condition. **C** Number of differentially accessible peaks detected using DESeq2, comparing naïve to 2h CpG stimulated pDCs, |FC|>2 and p<0.05. **D** Heatmap of normalized ATAC-Seq peak intensities (log2FC relative to the mean for each peak). Limited to peaks (16,607) that are condition-dependent with |FC|>2 and p<0.05 for at least one pairwise comparison of interest. **E** Differential motif analysis for cluster I and II from (**D**) using MEME Centrimo and the HOCOMOCO v11 motif database. Significant motifs were categorized into known TF families for visualization and interpretation.

### TFs show activation-dependent expression and chromatin accessibility

As shown above, pDC activation results in significant alterations of the chromatin landscape in pDCs making the DNA more or less accessible to specific TF families on a global level. We next analysed the impact of pDC activation on regions associated with TF genes themselves by evaluating regions ranging from 1kb upstream of the transcriptional start site (TSS) to 1kb downstream of the poly adenylation site. pDC activation altered the chromatin landscape of ~750 accessible regions associated with TF genes (|FC|>2, p<0.05, **Fig. 5A**). In detail, 2h stimulation of pDCs resulted in 627 peaks with increased accessibility and 126 peaks with decreased accessibility to regions associated with TF genes (**Fig. 5A**). 83% of all CpG-dependent chromatin regions in 2h stimulated pDCs exhibited increased DNA accessibility as compared to naïve pDCs. This suggests that most of the chromatin landscape associated with TF genes is “turned on” rather than being “turned off” after CpG stimulation. Finally, an integrative approach using the RNA-Seq and ATAC-Seq data was conducted analysing the differential chromatin states of regions associated with differentially expressed TF genes. This revealed 540 TF regions out of the overall ~750 chromatin regions that are significantly associated with a differential RNA expression of the respective TF gene (**Fig. 5B**). Out of these chromatin peaks we found 209 unique TF genes being associated with the differentially opened chromatin regions. Thus, pDC activation modulates the chromatin of most genes in more than one region associated with the respective gene, as shown here for the NFκB family members *Nfkb1* and *Rela* (**Fig. 5B**). To identify potential novel players in pDC biology after cell activation, we integrated the results of our motif analysis, the RNA expression levels, and chromatin states for all TFs. We focused our search on factors that fulfil the following criteria after pDC stimulation: (i) increased gene expression, (ii) enhanced chromatin accessibility, and (iii) enriched TF DNA binding motif in the genomic regions that are more accessible. Mining our dataset, we found that TFs already known to be important in TLR9-mediated signaling such as IRF and NFκB TFs met the requirement as expected. Additionally, members of the AP-1 family such as ATF3 and JUN, which received little mention for pDC biology in literature so far, also fulfilled these criteria. The candidates of all three families exhibited a significantly increased mRNA expression 2h after pDC activation as compared to naïve pDCs. At 6h after stimulation, expression remained at the same level (Jun, Rela), increased further (Irf7) or decreased (Atf3, Nfkb1). After 12h pDC stimulation, expression remained at the same level (Irf7, Atf3) or even decreased (Jun, Nfkb1, Rela) (**Fig. 5C**). In line with an increased expression of the selected TFs 2h after cell activation as compared to the naïve state, we found an increased accessibility of chromatin in the proximal promoter region of the *Irf7, Jun, Atf3, Nfkb1*, and *Rela* genes. Two regions of the *Nfkb1* gene, one proximal and another distal from the TSS of the gene, indicated increased DNA accessibility after CpG stimulation at 2h as compared to the naïve condition. While *Atf3, Nfkb1* and *Rela* are characterized by single or a small number of open chromatin peaks, several peaks in the *Irf7* and *Jun* gene were found, both proximal and after the TSS in the intergenic region. Of note, the core structural elements regulating gene expression for the proximal promoter and the intergenic regions were well conserved between mouse and human for all newly identified candidates (top panels, **Fig. 5D**). The potential relevance of the AP-1 factors for pDC biology was further investigated by searching for the common AP-1 motif (TGA[G/C]TCA) (Risse et al., 1989) among all open chromatin regions associated with pDC driving TF genes (*Runx2, Tcf4, Spib, Irf8, Bcl11a*). Using the MEME-FIMO search tool, we found an AP-1 motif in the proximal promoter site of the *Tcf4* gene which encodes the E2-2 protein (**Fig. 5E**). As AP-1 has not been implicated so far in E2-2 gene regulation this finding warrants further investigation. In summary, we found that pDC activation mostly “turns on” TF genes resulting in significant expression changes along with more accessible DNA in promoter and or intergenic regions. Moreover, we newly identified the AP-1 family as a set of TFs associated with pDC activation.

**Fig. 5.**
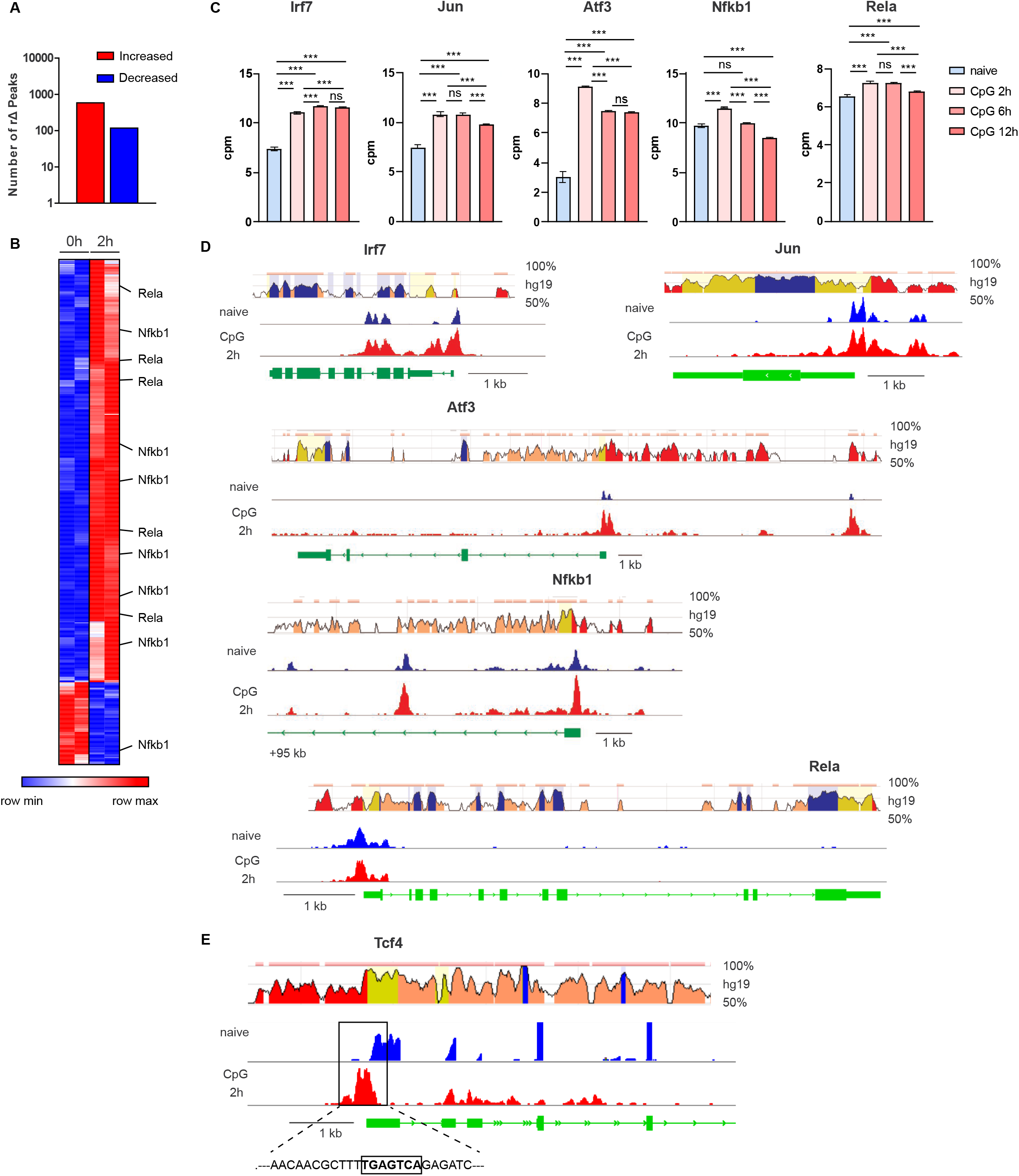
TFs show CpG-dependent expression and chromatin accessibility. **A** Number of differentially accessible peaks of genomic regions associated with TF genes detected using DESeq2 comparing naïve to 2h CpG stimulated pDCs, |FC|>2 and p<0.05. **B** Heatmap of normalized ATAC-Seq peak intensities (log2FC relative to the mean for each peak) limited to 540 peaks from (**A**) that are condition-dependent with |FC|>2 and p<0.05 for at least one pairwise comparison of interest. **C** The bar graph depicts normalized expression values obtained from RNA-Seq and statistics calculated with edgeR. **D, E** Top panel presents screen shots from the ECR (evolutionary conserved regions) Browser web site of the respective indicated gene. Exonic regions are shown in blue, intronic regions in pink, UTRs in yellow, and CNS in red. Bottom panels present ATAC-Seq peaks in naïve and CpG stimulated (2h) pDCs for the indicated genes visualized with IGV. The AP-1 motif within the promoter sequence of the *Tcf4* gene is highlighted in (**E**).

## Discussion

In this study we investigated the yet unknown global expression patterns of the TF reservoir of pDCs in in a time course after activation in combination with DNA accessibility analysis for implicated TF families. Combining RNA-Seq, ATAC-Seq, and GO analyses, we defined specific sets of TLR9-modulated TFs with known roles in pDC differentiation and function, but also TFs so far not implicated in pDC biology.

We used as the basis of our study the definition of the murine TF reservoir in the AnimalTFDB (Hu et al., 2019) and found that 70% of all genes annotated as TFs in the mouse genome (1,014 out of 1,636) were expressed in at least one condition, naïve or CpG-activated pDCs (2h, 6h, or 12h). These covered a wide range of TF classes defined by their respective DNA binding mechanisms. Interestingly, some TF families showed expression of all family members. Among those, we found factors that have been shown to be of particular importance in pDC biology, such as Runx2 of the Runt family (Sawai et al., 2013). Downstream GO analyses of RNA-Seq data allowed a biological classification of all TFs showing involvement in a wide variety of biological processes, such as the NFκB and JAK-STAT signaling. It has been well established that the production of type I IFN by pDCs upon TLR9 activation depends on the canonical TLR9-Myd88-NFκB/IRF7 signaling pathway (Tomasello et al., 2018). In this regard, it has been reported that NFκB and cREL are key players in pDC differentiation and survival programs after TLR9 activation by CpG. *Nfkb1*^-/-^ *cRel*^-/-^ double knock-out pDCs were still able to produce type I IFN upon CpG administration but failed to produce IL-6 or IL-12 and did not acquire a dendritic phenotype but rather underwent apoptosis (O’Keeffe et al., 2005). Here, we show for the first time the time-dependent patterns of gene expression for TFs involved in NFκB and JAK-STAT signaling upon pDC stimulation. Not only expression of these factors was enhanced in pDCs after CpG treatment, but also DNA binding sites for factors from the NFκB and JAK-STAT signaling pathways were identified as globally enriched in a differential motif analysis comparing regions with increased vs decreased chromatin accessibility. In addition, we found changed expression patterns of TFs important for circadian gene regulation in activated pDCs over time. In this regard, it has been reported that up to 10% of the transcriptome is under circadian regulation (Panda et al., 2002; Storch et al., 2002), suggesting that some pDC activation-dependent changes in gene expression may be under circadian control of global TF expression. Along this line, Silver et al. showed that TLR9 function is controlled by the circadian molecular clock in a number of cell types including DCs (Silver et al., 2012). Another group of TFs that show significant changes in expression after pDC activation could be classified as SMAD proteins, classical effectors of TGFβ signaling. It is known that stimulating DC progenitors with TGFβ accelerates DC differentiation, directing development toward cDCs (Felker et al., 2010). Also, one of the SMAD proteins, SMAD3, has been determined as a key player in determining cDC versus pDC cell fates (Jeong-Hwan Yoon, 2019). Interaction of SMAD proteins with known pDC driving factors such as Zeb2 have also been described (Vandewalle et al., 2009; Wu et al., 2016). Other SMAD members do not affect pDC numbers, as shown *in vivo* in *Smad7*-deficient mice (Lukas et al., 2017). Further, TFs involved in various processes of the endoplasmic reticulum are differentially expressed in TLR9 activated pDCs. Notably, mouse and human pDCs are morphologically characterized by an extensive rough ER, enabling them to rapidly secrete copious amounts of type I IFN after TLR7 and TLR9 stimulation (Alculumbre et al., 2018; Fitzgerald-Bocarsly et al., 2008). The enrichment of TFs involved in mRNA binding processes, sumoylation and epigenetic modifications further highlights the changing biology of pDCs in protein production, posttranslational protein modifications, and alteration of the physical DNA structure that regulates gene expression after cell activation. We hereby define a novel set of expressed TFs in TLR9 activated pDCs, thus identifying TFs involved in particular biological processes that may require further investigation for their functional role in activated pDCs. The global transcriptomics approach allows a comparison for the expression patterns of several TFs belonging to the same TF family or involved in the same biological process, which may help to further narrow down interesting candidates.

Using CpG as an optimal TLR9 agonist and focusing on early events after virus infection, we found that after pDC activation more of the pDC chromatin landscape is “turned on” rather than “turned off”, both globally in the genome and also among the regions associated with TF genes themselves. Specifically, about 80% of all regions that show significant chromatin changes exhibited increased accessibility for TFs. However, with regard to gene expression, 2h after pDC activation more genes were down-regulated than up-regulated as compared to the naïve state. One explanation could be that while DNA is more accessible, the TFs that possibly bind to these DNA stretches may inhibit rather than activate gene expression. An extensive motif analysis revealed that TFs belonging to the JAK-STAT and the NFκB signaling pathways exhibit increased accessibility to DNA binding regions after pDC stimulation. This underlines the importance of the JAK-STAT and NFκB signaling pathways in activated pDCs.

In contrast, Ets family members belonging to the Helix-turn-helix family of TFs and Zinc-coordinating zf-C2H2 TFs were both found to have less access to DNA after pDC activation. Ets family members include SPI1, also known as PU.1, which has been shown to drive the development of precursor cells toward cDC rather than pDC development (Chopin et al., 2019). Regarding pDC-driving cell fate TFs, IRF8 and RUNX2 belonging to the helix-turn-helix and β-scaffold TF groups, respectively, show motif enrichment in two sets of regions exhibiting increased versus decreased chromatin accessibility after pDC activation. Hence, cell fate TFs that drive pDC development both gain and lose access to distinct DNA regions after TLR9 activation.

Gene expression of the key pDC cell fate TFs IRF8, E2-2, and RUNX2 has been shown to steadily increase in expression during pDC precursor development into fully differentiated pDCs (Bornstein et al., 2014; Sasaki et al., 2012; Sawai et al., 2013; Tamura et al., 2005; Tsujimura et al., 2002). However, the role of these TFs for pDC survival and differentiation has not been investigated in detail after TLR9 activation. Here we observed different gene expression patterns for E2-2, and RUNX2 after pDC activation. E2-2 expression is strongly up-regulated at 2h and 6h of CpG stimulation vs no stimulation, but not at 12h after CpG activation vs steady state. Runx2, on the other hand, is strongly down-regulated at each CpG stimulation time point as compared to the naïve state.

Our results therefore warrant further investigations of pDC cell fate TFs to explore the biological relevance of distinct expression patterns as well as the simultaneous gain and loss of accessibility to DNA by modulation of chromatin after pDC activation. We found that IRF7, NFκB1, and RELA as well as ATF3 and JUN, two AP-1 family members, fulfil three criteria relevant in this context: They exhibit (i) increased gene expression, (ii) enhanced chromatin accessibility for their gene regions, and (iii) enriched TF DNA binding motifs in the accessible genomic regions after pDC stimulation. We used this integrative omics approach to identify potential novel players important in pDC biology after cell activation. While the role for IRF7, NFκB1, and RELA have been described in activated pDCs, there is little known about any function of AP-1 factors in pDCs. Activator Protein-1 (AP-1) was one of the first TFs to be described in the 1980s (Angel et al., 1987). It consists of a dimeric protein complex with members from the JUN, FOS, ATF, BATF, or MAF protein families (Eferl and Wagner, 2003; Shaulian and Karin, 2002). A shared feature between the members is a basic leucine-zipper (bZIP) domain which is required for dimerization and DNA binding. The AP-1 family of TFs are known to regulate various biological processes such as proliferation, differentiation, and cell survival (Eferl and Wagner, 2003; Murphy et al., 2013; Sopel et al., 2016; Wagner and Eferl, 2005). They have further been implicated in a variety of pathologies ranging from cardiovascular disease to cancer, hepatitis, and Parkinson’s disease (Meijer et al., 2012; Muslin, 2008; Uchihashi et al., 2011). A connection has been established between NFκB and AP-1 activity, which may be regulated by NFκB (Fujioka et al., 2004) suggesting a possible common molecular mechanism of these TFs in activated pDCs. Further, AP-1 has been shown to be required for spontaneous type I IFN production in pDCs, whereas type I IFN production triggered by pathogen receptor recognition such as TLR stimulation was not affected by AP-1 inhibition (Kim et al., 2014). In contrast, our *in silico* analyses suggest a close link between AP-1 factors and pDC biology after TLR9 stimulation: The AP-1 motif is present within the open chromatin region of the proximal promoter site of the *Tcf4* gene, a prominent pDC cell fate TF. Grajkowska *et al*. showed that there are two *Tcf4* isoforms, the expression of which is controlled during pDC differentiation by two respective promoters as well as distal enhancer regions within 600-900 kb 5’ and ~150 kb 3’ of the *Tcf4* gene (Grajkowska et al., 2017). However, the binding site of specific TFs to these cis-regulatory sites has not been fully evaluated. This calls for further investigations on the AP-1 binding site in activated pDCs newly identified in our study. One of the key AP-1 candidates in our investigation, ATF3, has been described as a negative regulator of antiviral signaling in Japanese encephalitis virus infection in mouse neuronal cells (Sood et al., 2017). The hallmark of pDCs is their importance in antiviral immune responses, pointing toward ATF3 as an interesting candidate to investigate in TLR9 activated pDCs. Another AP-1 family member, JUN, was the first oncogene to be described (Curran and Franza, 1988) and has since been studied in detail in the context of various tumor entities. In contrast, knowledge about its role in the context of infection is limited. For example, it has been shown to have a regulatory role in H5N1 influenza virus replication and host inflammation in mice (Xie et al., 2014). Our analyses revealed a distinct regulation of *Jun* expression and chromatin structure combined with an increased global DNA binding accessibility in pDCs after activation. Further studies are required to assess the role of *Jun* regulation in pDCs upon a microbial stimulus or in a chronically activated state that might unravel unknown functions of this TF in immunity. While targeting TFs for therapeutic purpose has been proven difficult so far, recent advances have been made through novel chemistries and the use of staples peptides to disrupt protein-protein interactions (Ball et al., 2016; Rezaei Araghi et al., 2018). Thus, the *in silico* analyses of the global TF reservoir in pDCs from our study led to the identification of novel candidates that warrant further investigation regarding their role in pDC biology, in particular after cell activation, which may lead to the development of novel therapeutics to treat infection, autoimmune disease and cancer.

## Author Contributions

RM analysed the data. SA performed BM Flt3-L pDC cultures and FACS sorted pDCs for the RNA-Seq and ATAC-Seq assays. PP and KK conducted RNA-Seq including primary analyses. RM, JA, and SS wrote the manuscript.

## Materials and Methods

### Mice

C57BL/6N mice were housed under specific pathogen-free conditions in the animal research facility of the University of Düsseldorf according to German animal welfare guidelines. All experiments were performed with sex and age matched littermates between 7 to 14 weeks of age.

### Generation and stimulation of BM-derived pDCs for RNA-Seq and ATAC-Seq

BM-derived Flt3-L cultured pDCs were generated as previously described (Scheu et al., 2008). For RNA-Seq, BM-derived pDCs (CD3^-^CD19^-^CD11c^+^CD11b^low^B220^+^SiglecH^+^ CD317^+^) were FACS purified using FACS Aria III (BD). The pDCs were left untreated or stimulated with 1μM CpG 2216 (Tib Molbiol, Nr. 930507l) complexed to transfection reagent DOTAP (Roche) for 2h, 6h or 12 h. RNA was isolated by using the NucleoSpin II RNA mini kit (Macherey-Nagel) and subjected to RNA-Seq. For ATAC-Seq BM-derived pDCs (CD3^-^CD19^-^CD11c^+^CD11b^low^B220^+^SiglecH^+^CD317^+^) were FACS purified using FACS Aria III (BD). The pDCs were left untreated or stimulated with 1μM CpG 2216 complexed to transfection reagent DOTAP (Roche) for 2h. At the end of stimulation time, cells were kept on ice and stained for 7AAD (BD). Live cells (7AAD^-^) were further purified by FACS and kept frozen in complete RPMI medium containing 5% DMSO. The frozen cells were transported on dry ice to Active Motif (Belgium) for ATAC-Seq.

The following antibodies have been used: CD3-PerCP (BD Bioscience, Clone: 145-2C11), CD19-PerCP-Cy5.5 (BD Bioscience, Clone:1D3), CD11c-PE-Cy7 (BioLegend, Clone: N418), CD11b-APC-Cy7 (BD Bioscience, Clone: M1/70), B220-FITC (BD Bioscience, Clone: RA3-6B2), SiglecH-APC (BioLegend, Clone 551), CD317-PE (eBioscience/Thermoscientific, Clone: ebio927).

### RNA-Seq Analyses

DNase digested total RNA samples used for transcriptome analyses were quantified (Qubit RNA HS Assay, Thermo Fisher Scientific) and quality measured by capillary electrophoresis using the Fragment Analyzer and the ‘Total RNA Standard Sensitivity Assay’ (Agilent Technologies, Inc. Santa Clara, USA). All samples in this study showed high RNA Quality Numbers (RQN; mean = 9.9). The library preparation was performed according to the manufacturer’s protocol using the Illumina^®^ ‘TruSeq Stranded mRNA Library Prep Kit’. Briefly, 200 ng total RNA were used for mRNA capturing, fragmentation, the synthesis of cDNA, adapter ligation and library amplification. Bead purified libraries were normalized and sequenced on the HiSeq 3000/4000 system (Illumina Inc. San Diego, USA) with a read setup of SR 1×150 bp. The bcl2fastq tool was used to convert the bcl files to fastq files as well for adapter trimming and demultiplexing.

Data analyses on fastq files were conducted with CLC Genomics Workbench (version 11.0.1, QIAGEN, Venlo. NL). The reads of all probes were adapter trimmed (Illumina TruSeq) and quality trimmed (using the default parameters: bases below Q13 were trimmed from the end of the reads, ambiguous nucleotides maximal 2). Mapping was done against the *Mus musculus* (mm10; GRCm38.86) (March 24, 2017) genome sequence. Samples (three biological replicates each) were grouped according to their respective experimental condition. Raw counts were next re-uploaded to the Galaxy web platform. The public server at usegalaxy.org was used to perform multi-group comparisons (Afgan et al., 2016). Differential expression of genes between any two conditions was calculated using the edgeR quasi-likelihood pipeline which uses negative binomial generalized linear models with F-test (Liu et al., 2015; Robinson et al., 2010). Low expressing genes were filtered with a count-per-million (CPM) value cut-off that was calculated based on the average library size of our RNA-Seq experiment (Chen et al., 2016). The resulting p values were corrected for multiple testing by the false discovery rate (FDR) (Benjamini, 1995). A p value of <0.05 was considered significant. RNA-Seq data are deposited with NCBI’s Gene Expression Omnibus (GEO) and are accessible through GEO Series accession number GSE170750 (https://www.ncbi.nlm.nih.gov/geo/query/acc.cgi?acc=GSE170750).

### ATAC-Seq

Cells were harvested and frozen in culture media containing FBS and 5% DMSO. Cryopreserved cells were sent to Active Motif to perform the ATAC-Seq assay. The cells were then thawed in a 37°C water bath, pelleted, washed with cold PBS, and tagmented as previously described (Buenrostro et al., 2013), with some modifications (Corces et al., 2017). Briefly, cell pellets were resuspended in lysis buffer, pelleted, and tagmented using the enzyme and buffer provided in the Nextera Library Prep Kit (Illumina). Tagmented DNA was then purified using the MinElute PCR purification kit (Qiagen), amplified with 10 cycles of PCR, and purified using Agencourt AMPure SPRI beads (Beckman Coulter). Resulting material was quantified using the KAPA Library Quantification Kit for Illumina platforms (KAPA Biosystems), and sequenced with PE42 sequencing on the NextSeq 500 sequencer (Illumina).

Reads were aligned using the BWA algorithm (mem mode; default settings). Duplicate reads were removed, only reads mapping as matched pairs and only uniquely mapped reads (mapping quality ≥1) were used for further analysis. Alignments were extended *in silico* at their 3’-ends to a length of 200 bp and assigned to 32-nt bins along the genome. The resulting histograms (genomic “signal maps”) were stored in bigWig files. Peaks were identified using the MACS 2.1.0 algorithm at a cut off of p-value 1e-7, without control file, and with the –nomodel option. Peaks that were on the ENCODE blacklist of known false ATAC-Seq peaks were removed. Signal maps and peak locations were used as input data to Active Motifs proprietary analysis program, which creates Excel tables containing detailed information on sample comparison, peak metrics, peak locations, and gene annotations. For differential analysis, reads were counted in all merged peak regions (using Subread), and the replicates for each condition were compared using DESeq2. ATAC-Seq data are deposited with NCBI’s GEO and are accessible through GEO Series accession number GSE171075 (https://www.ncbi.nlm.nih.gov/geo/query/acc.cgi?acc=GSE171075).

### Downstream analyses and visualization of omics data

Volcano plots were created using ggplot2 (Wickham, 2016) and ggrepel (Slowikowski, 2020). Heatmaps were created using Morpheus (https://software.broadinstitute.org/morpheus). Pearson correlation matrices were calculated in R and plotted as heatmaps using gplots (Gregory R. Warnes, 2020). Pathway analyses for different gene ontology (GO) terms and subsequent functional classification and annotation clustering were performed using the Database for Annotation, Visualization and Integrated Discovery (DAVID) (Huang da et al., 2009). Evolutionary conserved regions (ECR) for selected genes were shown by taking a screenshot from the ECR browser (Ovcharenko et al., 2004). Bar graphs were plotted in Gradphpad Prism version 8.4.3 on Windows (GraphPad Software, La Jolla California USA, www.graphpad.com). ATAC-Seq peaks were visualized using IGV (Robinson et al., 2011; Thorvaldsdottir et al., 2013).

### TF Motif Analyses

ATAC-Seq regions that indicated differentially accessible chromatin regions between naive and 2h CpG stimulated samples (DESeq2, |FC|>2, p<0.05) were used for motif analysis. The regions were adjusted to the same size (500bp). The MEME-Centrimo differential motif analysis pipeline (Bailey and Machanick, 2012) was run on the fasta files representing each chromatin region (significantly increased vs decreased chromatin access after CpG stimulation) to identify overrepresented motifs, using default parameters and the HOCOMOCO v11 motif database. The search for the AP-1 motif among selected sequences was performed with MEME-FIMO.

## Acknowledgements

This work was funded by the German Research Foundation (DFG – SCHE692/6-1) and the Manchot Graduate Schools ‘Molecules of Infection III’ to SS and the DFG EXC 1003, Grant FF-2014-01 Cells in Motion–Cluster of Excellence, Münster, Germany, and the DFG FOR2107 AL1145/5-2 to JA. Computational support of the Zentrum für Informations- und Medientechnologie, especially the HPC team (High Performance Computing) at the University of Düsseldorf is acknowledged. We thank Johannes Ptok and Heiner Schaal (Institute of Virology, University of Düsseldorf) for critical reading of the manuscript.

---

**SUPPLEMENTAL TABLE S1.**
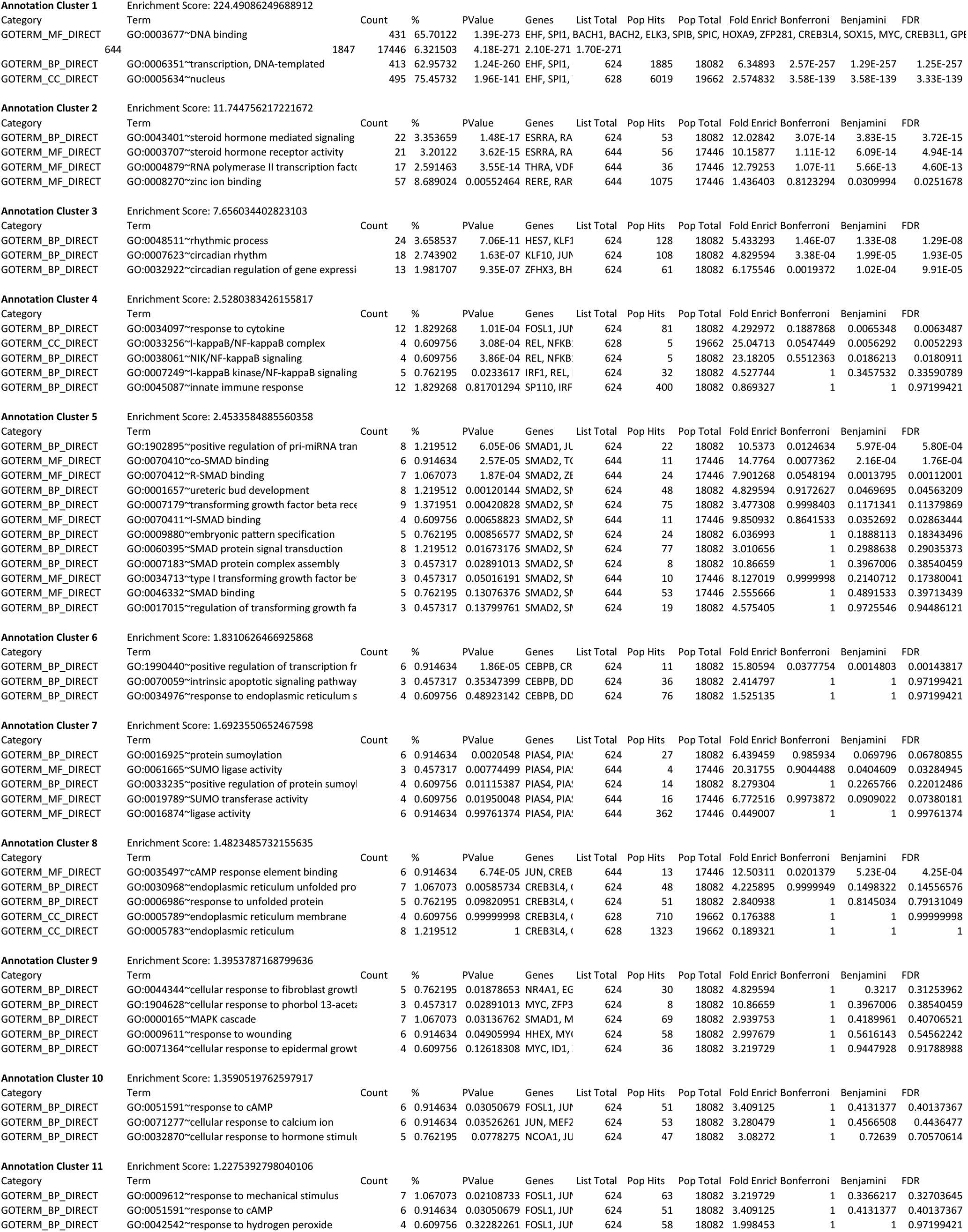
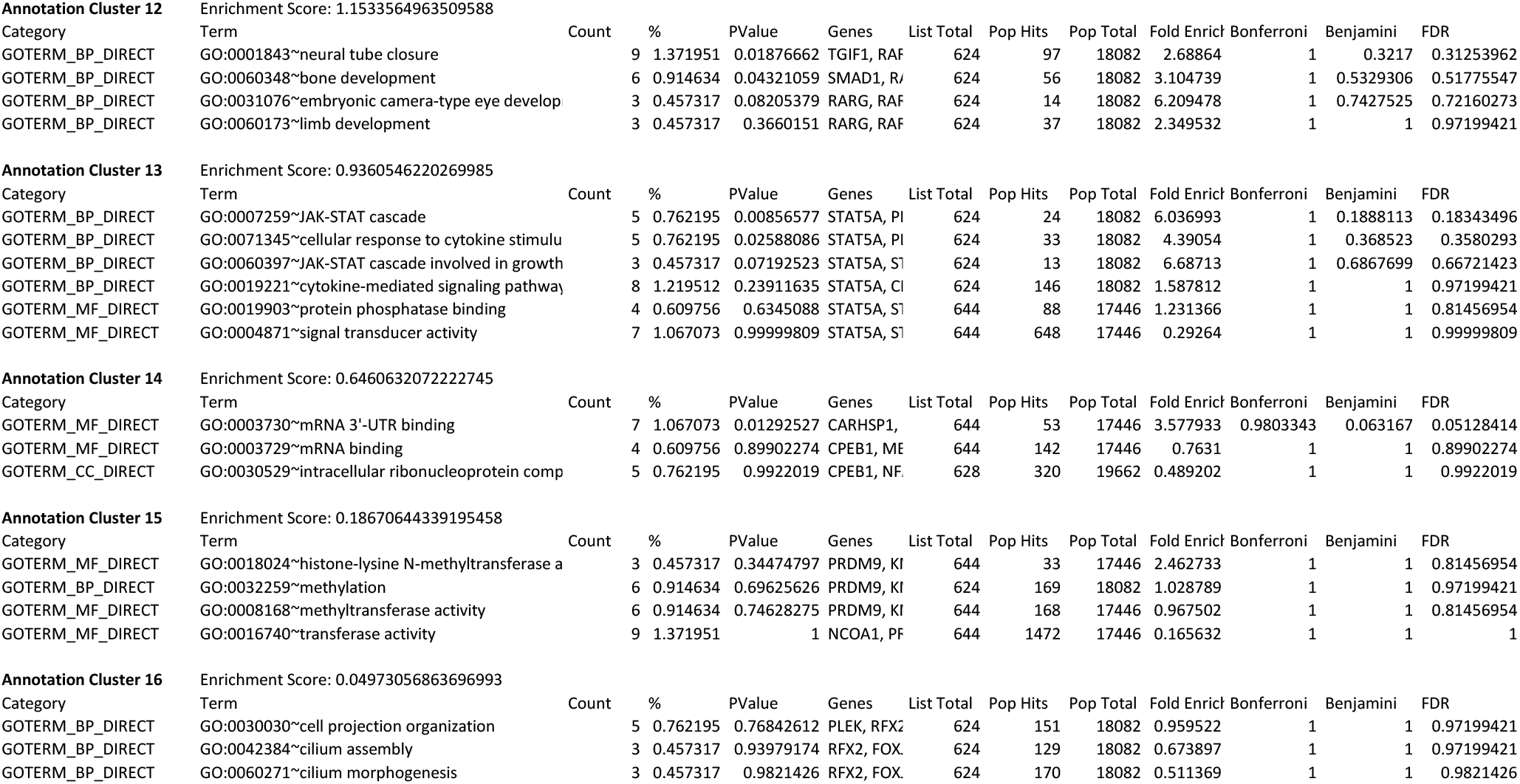
Functional cluster analysis with 661 CpG-dependent TF genes

**SUPPLEMENTAL TABLE S2.**
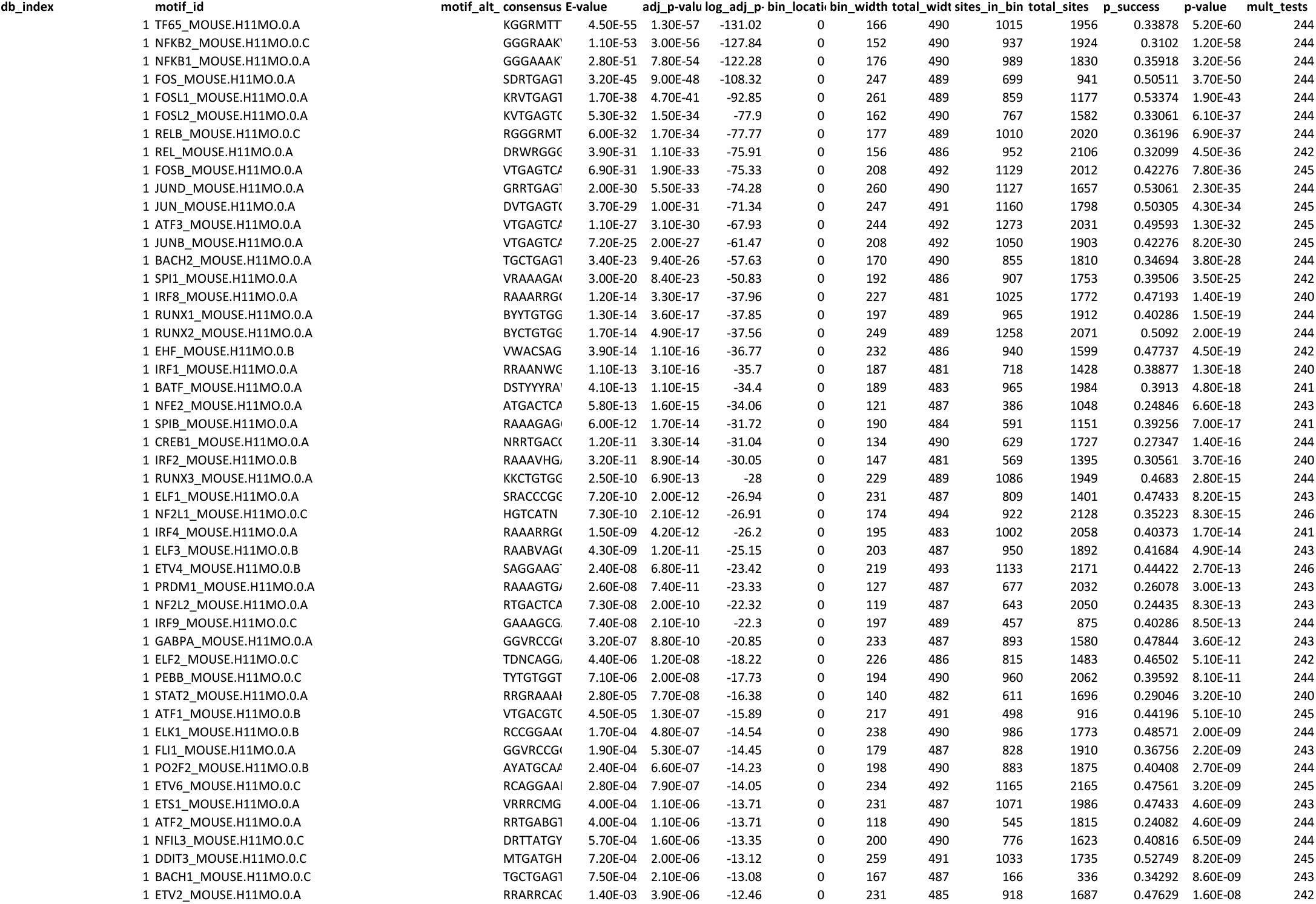
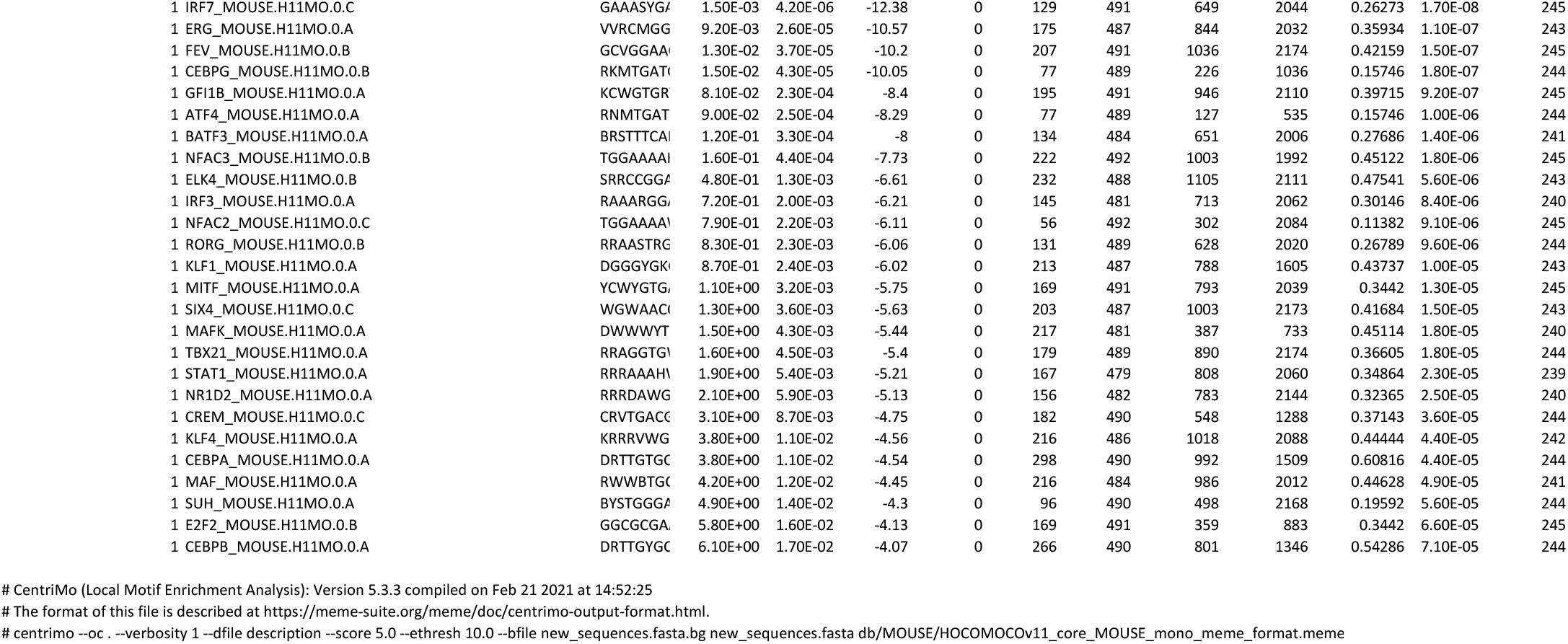
Centrimo motif enrichment analysis using the HOCOMOCO database and 2174 gene promoter associated regions with increased chromatin accessibility after pDC activation

## References

Afgan, E., Baker, D., van den Beek, M., Blankenberg, D., Bouvier, D., Cech, M., Chilton, J., Clements, D., Coraor, N., Eberhard, C., et al. (2016). The Galaxy platform for accessible, reproducible and collaborative biomedical analyses: 2016 update. Nucleic Acids Res 44, W3–W10.

Alculumbre, S.G., Saint-Andre, V., Di Domizio, J., Vargas, P., Sirven, P., Bost, P., Maurin, M., Maiuri, P., Wery, M., Roman, M.S., et al. (2018). Diversification of human plasmacytoid predendritic cells in response to a single stimulus. Nat Immunol 19, 63–75.

Ali, S., Mann-Nuttel, R., Schulze, A., Richter, L., Alferink, J., and Scheu, S. (2019). Sources of Type I Interferons in Infectious Immunity: Plasmacytoid Dendritic Cells Not Always in the Driver’s Seat. Front Immunol 10, 778.

Angel, P., Imagawa, M., Chiu, R., Stein, B., Imbra, R.J., Rahmsdorf, H.J., Jonat, C., Herrlich, P., and Karin, M. (1987). Phorbol ester-inducible genes contain a common cis element recognized by a TPA-modulated trans-acting factor. Cell 49, 729–739.

Aravind, L., Anantharaman, V., Balaji, S., Babu, M.M., and Iyer, L.M. (2005). The many faces of the helix-turn-helix domain: transcription regulation and beyond. FEMS Microbiol Rev 29, 231–262.

Asselin-Paturel, C., Boonstra, A., Dalod, M., Durand, I., Yessaad, N., Dezutter-Dambuyant, C., Vicari, A., O’Garra, A., Biron, C., Briere, F., and Trinchieri, G. (2001). Mouse type I IFN-producing cells are immature APCs with plasmacytoid morphology. Nat Immunol 2, 1144–1150.

Bailey, T.L., and Machanick, P. (2012). Inferring direct DNA binding from ChIP-seq. Nucleic Acids Res 40, e128.

Ball, D.P., Lewis, A.M., Williams, D., Resetca, D., Wilson, D.J., and Gunning, P.T. (2016). Signal transducer and activator of transcription 3 (STAT3) inhibitor, S3I-201, acts as a potent and non-selective alkylating agent. Oncotarget 7, 20669–20679.

Bauer, J., Dress, R.J., Schulze, A., Dresing, P., Ali, S., Deenen, R., Alferink, J., and Scheu, S. (2016). Cutting Edge: IFN-beta Expression in the Spleen Is Restricted to a Subpopulation of Plasmacytoid Dendritic Cells Exhibiting a Specific Immune Modulatory Transcriptome Signature. J Immunol 196, 4447–4451.

Benjamini, Y.H., Yosef (1995). Controlling the false discovery rate: a practical and powerful approach to multiple testing. Journal of the Royal Statistical Society, Series B 57 (1): 289–300.

Bornstein, C., Winter, D., Barnett-Itzhaki, Z., David, E., Kadri, S., Garber, M., and Amit, I. (2014). A negative feedback loop of transcription factors specifies alternative dendritic cell chromatin States. Mol Cell 56, 749–762.

Buenrostro, J.D., Giresi, P.G., Zaba, L.C., Chang, H.Y., and Greenleaf, W.J. (2013). Transposition of native chromatin for fast and sensitive epigenomic profiling of open chromatin, DNA-binding proteins and nucleosome position. Nat Methods 10, 1213–1218.

Chauvistre, H., and Sere, K. (2020). Epigenetic aspects of DC development and differentiation. Mol Immunol 128, 116–124.

Chen, Y., Lun, A.T., and Smyth, G.K. (2016). From reads to genes to pathways: differential expression analysis of RNA-Seq experiments using Rsubread and the edgeR quasi-likelihood pipeline. F1000Res 5, 1438.

Chopin, M., Lun, A.T., Zhan, Y., Schreuder, J., Coughlan, H., D’Amico, A., Mielke, L.A., Almeida, F.F., Kueh, A.J., Dickins, R.A., et al. (2019). Transcription Factor PU.1 Promotes Conventional Dendritic Cell Identity and Function via Induction of Transcriptional Regulator DC-SCRIPT. Immunity 50, 77–90 e75.

Chopin, M., Preston, S.P., Lun, A.T.L., Tellier, J., Smyth, G.K., Pellegrini, M., Belz, G.T., Corcoran, L.M., Visvader, J.E., Wu, L., and Nutt, S.L. (2016). RUNX2 Mediates Plasmacytoid Dendritic Cell Egress from the Bone Marrow and Controls Viral Immunity. Cell Rep 15, 866–878.

Christensen, S.R., Shupe, J., Nickerson, K., Kashgarian, M., Flavell, R.A., and Shlomchik, M.J. (2006). Toll-like receptor 7 and TLR9 dictate autoantibody specificity and have opposing inflammatory and regulatory roles in a murine model of lupus. Immunity 25, 417–428.

Cisse, B., Caton, M.L., Lehner, M., Maeda, T., Scheu, S., Locksley, R., Holmberg, D., Zweier, C., den Hollander, N.S., Kant, S.G., et al. (2008). Transcription factor E2-2 is an essential and specific regulator of plasmacytoid dendritic cell development. Cell 135, 37–48.

Corces, M.R., Trevino, A.E., Hamilton, E.G., Greenside, P.G., Sinnott-Armstrong, N.A., Vesuna, S., Satpathy, A.T., Rubin, A.J., Montine, K.S., Wu, B., et al. (2017). An improved ATAC-seq protocol reduces background and enables interrogation of frozen tissues. Nat Methods 14, 959–962.

Curran, T., and Franza, B.R., Jr. (1988). Fos and Jun: the AP-1 connection. Cell 55, 395–397.

Deaton, A.M., and Bird, A. (2011). CpG islands and the regulation of transcription. Genes Dev 25, 1010–1022.

Decque, A., Joffre, O., Magalhaes, J.G., Cossec, J.C., Blecher-Gonen, R., Lapaquette, P., Silvin, A., Manel, N., Joubert, P.E., Seeler, J.S., et al. (2016). Sumoylation coordinates the repression of inflammatory and anti-viral gene-expression programs during innate sensing. Nat Immunol 17, 140–149.

Diebold, S.S., Kaisho, T., Hemmi, H., Akira, S., and Reis e Sousa, C. (2004). Innate antiviral responses by means of TLR7-mediated recognition of single-stranded RNA. Science 303, 1529–1531.

Drouin, J. (2014). Minireview: pioneer transcription factors in cell fate specification. Mol Endocrinol 28, 989–998.

Eferl, R., and Wagner, E.F. (2003). AP-1: a double-edged sword in tumorigenesis. Nat Rev Cancer 3, 859–868.

Elkon, K.B., and Wiedeman, A. (2012). Type I IFN system in the development and manifestations of SLE. Curr Opin Rheumatol 24, 499–505.

Felker, P., Sere, K., Lin, Q., Becker, C., Hristov, M., Hieronymus, T., and Zenke, M. (2010). TGF-beta1 accelerates dendritic cell differentiation from common dendritic cell progenitors and directs subset specification toward conventional dendritic cells. J Immunol 185, 5326–5335.

Fitzgerald-Bocarsly, P., Dai, J., and Singh, S. (2008). Plasmacytoid dendritic cells and type I IFN: 50 years of convergent history. Cytokine Growth Factor Rev 19, 3–19.

Fujioka, S., Niu, J., Schmidt, C., Sclabas, G.M., Peng, B., Uwagawa, T., Li, Z., Evans, D.B., Abbruzzese, J.L., and Chiao, P.J. (2004). NF-kappaB and AP-1 connection: mechanism of NF-kappaB-dependent regulation of AP-1 activity. Mol Cell Biol 24, 7806–7819.

Fulton, D.L., Sundararajan, S., Badis, G., Hughes, T.R., Wasserman, W.W., Roach, J.C., and Sladek, R. (2009). TFCat: the curated catalog of mouse and human transcription factors. Genome Biol 10, R29.

Gilliet, M., Cao, W., and Liu, Y.J. (2008). Plasmacytoid dendritic cells: sensing nucleic acids in viral infection and autoimmune diseases. Nat Rev Immunol 8, 594–606.

Grajkowska, L.T., Ceribelli, M., Lau, C.M., Warren, M.E., Tiniakou, I., Nakandakari Higa, S., Bunin, A., Haecker, H., Mirny, L.A., Staudt, L.M., and Reizis, B. (2017). Isoform-Specific Expression and Feedback Regulation of E Protein TCF4 Control Dendritic Cell Lineage Specification. Immunity 46, 65–77.

Gregory R. Warnes, B.B., Lodewijk Bonebakker, Robert Gentleman, Wolfgang Huber, Andy Liaw, Thomas Lumley, Martin Maechler, Arni Magnusson, Steffen Moeller, Marc Schwartz and Bill Venables (2020). gplots: Various R Programming Tools for Plotting Data. ScienceOpen.

Hayashi, T., Beck, L., Rossetto, C., Gong, X., Takikawa, O., Takabayashi, K., Broide, D.H., Carson, D.A., and Raz, E. (2004). Inhibition of experimental asthma by indoleamine 2,3-dioxygenase. J Clin Invest 114, 270–279.

Hornung, V., Rothenfusser, S., Britsch, S., Krug, A., Jahrsdorfer, B., Giese, T., Endres, S., and Hartmann, G. (2002). Quantitative expression of toll-like receptor 1-10 mRNA in cellular subsets of human peripheral blood mononuclear cells and sensitivity to CpG oligodeoxynucleotides. J Immunol 168, 4531–4537.

Hu, H., Miao, Y.R., Jia, L.H., Yu, Q.Y., Zhang, Q., and Guo, A.Y. (2019). AnimalTFDB 3.0: a comprehensive resource for annotation and prediction of animal transcription factors. Nucleic Acids Res 47, D33–D38.

Huang da, W., Sherman, B.T., and Lempicki, R.A. (2009). Systematic and integrative analysis of large gene lists using DAVID bioinformatics resources. Nat Protoc 4, 44–57.

Ishii, K.J., and Akira, S. (2006). Innate immune recognition of, and regulation by, DNA. Trends Immunol 27, 525–532.

Jeong-Hwan Yoon, E.B., Katsuko Sudo, Jin Soo Han, Seok Hee Park, Susumu Nakae, Tadashi Yamashita, In-Kyu Lee, Ji Hyeon Ju, Isao Matsumoto, Takayuki Sumida, Masahiko Kuroda, Keiji Miyazawa, Mitsuyasu Kato, Mizuko Mamura (2019). SMAD3 Determines Conventional versus Plasmacytoid Dendritic Cell Fates. bioRxiv.

Kanamori, M., Konno, H., Osato, N., Kawai, J., Hayashizaki, Y., and Suzuki, H. (2004). A genome-wide and nonredundant mouse transcription factor database. Biochem Biophys Res Commun 322, 787–793.

Kim, S., Kaiser, V., Beier, E., Bechheim, M., Guenthner-Biller, M., Ablasser, A., Berger, M., Endres, S., Hartmann, G., and Hornung, V. (2014). Self-priming determines high type I IFN production by plasmacytoid dendritic cells. Eur J Immunol 44, 807–818.

Kulakovskiy, I.V., Vorontsov, I.E., Yevshin, I.S., Sharipov, R.N., Fedorova, A.D., Rumynskiy, E.I., Medvedeva, Y.A., Magana-Mora, A., Bajic, V.B., Papatsenko, D.A., et al. (2018). HOCOMOCO: towards a complete collection of transcription factor binding models for human and mouse via large-scale ChIP-Seq analysis. Nucleic Acids Res 46, D252–D259.

Le Mercier, I., Poujol, D., Sanlaville, A., Sisirak, V., Gobert, M., Durand, I., Dubois, B., Treilleux, I., Marvel, J., Vlach, J., et al. (2013). Tumor promotion by intratumoral plasmacytoid dendritic cells is reversed by TLR7 ligand treatment. Cancer Res 73, 4629–4640.

Lee, J.U., Kim, L.K., and Choi, J.M. (2018). Revisiting the Concept of Targeting NFAT to Control T Cell Immunity and Autoimmune Diseases. Front Immunol 9, 2747.

Li, S., Wu, J., Zhu, S., Liu, Y.J., and Chen, J. (2017). Disease-Associated Plasmacytoid Dendritic Cells. Front Immunol 8, 1268.

Libermann, T.A., and Zerbini, L.F. (2006). Targeting transcription factors for cancer gene therapy. Curr Gene Ther 6, 17–33.

Lin, Q., Chauvistre, H., Costa, I.G., Gusmao, E.G., Mitzka, S., Hanzelmann, S., Baying, B., Klisch, T., Moriggl, R., Hennuy, B., et al. (2015). Epigenetic program and transcription factor circuitry of dendritic cell development. Nucleic Acids Res 43, 9680–9693.

Liu, R., Holik, A.Z., Su, S., Jansz, N., Chen, K., Leong, H.S., Blewitt, M.E., Asselin-Labat, M.L., Smyth, G.K., and Ritchie, M.E. (2015). Why weight? Modelling sample and observational level variability improves power in RNA-seq analyses. Nucleic Acids Res 43, e97.

Liu, Y.J. (2005). IPC: professional type 1 interferon-producing cells and plasmacytoid dendritic cell precursors. Annu Rev Immunol 23, 275–306.

Lou, Y., Liu, C., Lizee, G., Peng, W., Xu, C., Ye, Y., Rabinovich, B.A., Hailemichael, Y., Gelbard, A., Zhou, D., et al. (2011). Antitumor activity mediated by CpG: the route of administration is critical. J Immunother 34, 279–288.

Lukas, D., Yogev, N., Kel, J.M., Regen, T., Mufazalov, I.A., Tang, Y., Wanke, F., Reizis, B., Muller, W., Kurschus, F.C., et al. (2017). TGF-beta inhibitor Smad7 regulates dendritic cell-induced autoimmunity. Proc Natl Acad Sci U S A 114, E1480–E1489.

Meijer, C.A., Le Haen, P.A., van Dijk, R.A., Hira, M., Hamming, J.F., van Bockel, J.H., and Lindeman, J.H. (2012). Activator protein-1 (AP-1) signalling in human atherosclerosis: results of a systematic evaluation and intervention study. Clin Sci (Lond) 122, 421–428.

Murphy, T.L., Tussiwand, R., and Murphy, K.M. (2013). Specificity through cooperation: BATF-IRF interactions control immune-regulatory networks. Nat Rev Immunol 13, 499–509.

Muslin, A.J. (2008). MAPK signalling in cardiovascular health and disease: molecular mechanisms and therapeutic targets. Clin Sci (Lond) 115, 203–218.

O’Keeffe, M., Grumont, R.J., Hochrein, H., Fuchsberger, M., Gugasyan, R., Vremec, D., Shortman, K., and Gerondakis, S. (2005). Distinct roles for the NF-kappaB1 and c-Rel transcription factors in the differentiation and survival of plasmacytoid and conventional dendritic cells activated by TLR-9 signals. Blood 106, 3457–3464.

Ovcharenko, I., Nobrega, M.A., Loots, G.G., and Stubbs, L. (2004). ECR Browser: a tool for visualizing and accessing data from comparisons of multiple vertebrate genomes. Nucleic Acids Res 32, W280–286.

Panda, S., Antoch, M.P., Miller, B.H., Su, A.I., Schook, A.B., Straume, M., Schultz, P.G., Kay, S.A., Takahashi, J.S., and Hogenesch, J.B. (2002). Coordinated transcription of key pathways in the mouse by the circadian clock. Cell 109, 307–320.

Patel, L., Abate, C., and Curran, T. (1990). Altered protein conformation on DNA binding by Fos and Jun. Nature 347, 572–575.

Reizis, B. (2019). Plasmacytoid Dendritic Cells: Development, Regulation, and Function. Immunity 50, 37–50.

Rezaei Araghi, R., Bird, G.H., Ryan, J.A., Jenson, J.M., Godes, M., Pritz, J.R., Grant, R.A., Letai, A., Walensky, L.D., and Keating, A.E. (2018). Iterative optimization yields Mcl-1-targeting stapled peptides with selective cytotoxicity to Mcl-1-dependent cancer cells. Proc Natl Acad Sci U S A 115, E886–E895.

Risse, G., Jooss, K., Neuberg, M., Bruller, H.J., and Muller, R. (1989). Asymmetrical recognition of the palindromic AP1 binding site (TRE) by Fos protein complexes. EMBO J 8, 3825–3832.

Robinson, J.T., Thorvaldsdottir, H., Winckler, W., Guttman, M., Lander, E.S., Getz, G., and Mesirov, J.P. (2011). Integrative genomics viewer. Nat Biotechnol 29, 24–26.

Robinson, M.D., McCarthy, D.J., and Smyth, G.K. (2010). edgeR: a Bioconductor package for differential expression analysis of digital gene expression data. Bioinformatics 26, 139–140.

Saas, P., and Perruche, S. (2012). Functions of TGF-beta-exposed plasmacytoid dendritic cells. Crit Rev Immunol 32, 529–553.

Salio, M., Palmowski, M.J., Atzberger, A., Hermans, I.F., and Cerundolo, V. (2004). CpG-matured murine plasmacytoid dendritic cells are capable of in vivo priming of functional CD8 T cell responses to endogenous but not exogenous antigens. J Exp Med 199, 567–579.

Sasaki, I., Hoshino, K., Sugiyama, T., Yamazaki, C., Yano, T., Iizuka, A., Hemmi, H., Tanaka, T., Saito, M., Sugiyama, M., et al. (2012). Spi-B is critical for plasmacytoid dendritic cell function and development. Blood 120, 4733–4743.

Sawai, C.M., Sisirak, V., Ghosh, H.S., Hou, E.Z., Ceribelli, M., Staudt, L.M., and Reizis, B. (2013). Transcription factor Runx2 controls the development and migration of plasmacytoid dendritic cells. J Exp Med 210, 2151–2159.

Scheu, S., Dresing, P., and Locksley, R.M. (2008). Visualization of IFNbeta production by plasmacytoid versus conventional dendritic cells under specific stimulation conditions in vivo. Proc Natl Acad Sci U S A 105, 20416–20421.

Serebreni, L., and Stark, A. (2020). Insights into gene regulation: From regulatory genomic elements to DNA-protein and protein-protein interactions. Curr Opin Cell Biol 70, 58–66.

Shaulian, E., and Karin, M. (2002). AP-1 as a regulator of cell life and death. Nat Cell Biol 4, E131–136.

Silver, A.C., Arjona, A., Walker, W.E., and Fikrig, E. (2012). The circadian clock controls toll-like receptor 9-mediated innate and adaptive immunity. Immunity 36, 251–261.

Slowikowski, K. (2020). ggrepel: Automatically Position Non-Overlapping Text Labels with ‘ggplot2’.

Sood, V., Sharma, K.B., Gupta, V., Saha, D., Dhapola, P., Sharma, M., Sen, U., Kitajima, S., Chowdhury, S., Kalia, M., and Vrati, S. (2017). ATF3 negatively regulates cellular antiviral signaling and autophagy in the absence of type I interferons. Sci Rep 7, 8789.

Sopel, N., Graser, A., Mousset, S., and Finotto, S. (2016). The transcription factor BATF modulates cytokine-mediated responses in T cells. Cytokine Growth Factor Rev 30, 39–45.

Storch, K.F., Lipan, O., Leykin, I., Viswanathan, N., Davis, F.C., Wong, W.H., and Weitz, C.J. (2002). Extensive and divergent circadian gene expression in liver and heart. Nature 417, 78–83.

Swiecki, M., and Colonna, M. (2015). The multifaceted biology of plasmacytoid dendritic cells. Nat Rev Immunol 15, 471–485.

Takayanagi, H., Kim, S., Matsuo, K., Suzuki, H., Suzuki, T., Sato, K., Yokochi, T., Oda, H., Nakamura, K., Ida, N., et al. (2002). RANKL maintains bone homeostasis through c-Fos-dependent induction of interferon-beta. Nature 416, 744–749.

Tamura, T., Tailor, P., Yamaoka, K., Kong, H.J., Tsujimura, H., O’Shea, J.J., Singh, H., and Ozato, K. (2005). IFN regulatory factor-4 and −8 govern dendritic cell subset development and their functional diversity. J Immunol 174, 2573–2581.

Thorvaldsdottir, H., Robinson, J.T., and Mesirov, J.P. (2013). Integrative Genomics Viewer (IGV): high-performance genomics data visualization and exploration. Brief Bioinform 14, 178–192.

Tomasello, E., Naciri, K., Chelbi, R., Bessou, G., Fries, A., Gressier, E., Abbas, A., Pollet, E., Pierre, P., Lawrence, T., et al. (2018). Molecular dissection of plasmacytoid dendritic cell activation in vivo during a viral infection. EMBO J 37.

Trinchieri, G., and Santoli, D. (1978). Anti-viral activity induced by culturing lymphocytes with tumor-derived or virus-transformed cells. Enhancement of human natural killer cell activity by interferon and antagonistic inhibition of susceptibility of target cells to lysis. J Exp Med 147, 1314–1333.

Tsujimura, H., Nagamura-Inoue, T., Tamura, T., and Ozato, K. (2002). IFN consensus sequence binding protein/IFN regulatory factor-8 guides bone marrow progenitor cells toward the macrophage lineage. J Immunol 169, 1261–1269.

Uchihashi, S., Fukumoto, H., Onoda, M., Hayakawa, H., Ikushiro, S., and Sakaki, T. (2011). Metabolism of the c-Fos/activator protein-1 inhibitor T-5224 by multiple human UDP-glucuronosyltransferase isoforms. Drug Metab Dispos 39, 803–813.

Vandewalle, C., Van Roy, F., and Berx, G. (2009). The role of the ZEB family of transcription factors in development and disease. Cell Mol Life Sci 66, 773–787.

Vaquerizas, J.M., Kummerfeld, S.K., Teichmann, S.A., and Luscombe, N.M. (2009). A census of human transcription factors: function, expression and evolution. Nat Rev Genet 10, 252–263.

Wagner, E.F., and Eferl, R. (2005). Fos/AP-1 proteins in bone and the immune system. Immunol Rev 208, 126–140.

Weiss, M.A., Ellenberger, T., Wobbe, C.R., Lee, J.P., Harrison, S.C., and Struhl, K. (1990). Folding transition in the DNA-binding domain of GCN4 on specific binding to DNA. Nature 347, 575–578.

Wickham, H. (2016). ggplot2: Elegant Graphics for Data Analysis. Springer-Verlag New York.

Wingender, E., Schoeps, T., Haubrock, M., Krull, M., and Donitz, J. (2018). TFClass: expanding the classification of human transcription factors to their mammalian orthologs. Nucleic Acids Res 46, D343–D347.

Wu, J., Li, S., Li, T., Lv, X., Zhang, M., Zang, G., Qi, C., Liu, Y.J., Xu, L., and Chen, J. (2019). pDC Activation by TLR7/8 Ligand CL097 Compared to TLR7 Ligand IMQ or TLR9 Ligand CpG. J Immunol Res 2019, 1749803.

Wu, X., Briseno, C.G., Grajales-Reyes, G.E., Haldar, M., Iwata, A., Kretzer, N.M., Kc, W., Tussiwand, R., Higashi, Y., Murphy, T.L., and Murphy, K.M. (2016). Transcription factor Zeb2 regulates commitment to plasmacytoid dendritic cell and monocyte fate. Proc Natl Acad Sci U S A 113, 14775–14780.

Xie, J., Zhang, S., Hu, Y., Li, D., Cui, J., Xue, J., Zhang, G., Khachigian, L.M., Wong, J., Sun, L., and Wang, M. (2014). Regulatory roles of c-jun in H5N1 influenza virus replication and host inflammation. Biochim Biophys Acta 1842, 2479–2488.

Zhang, H.M., Chen, H., Liu, W., Liu, H., Gong, J., Wang, H., and Guo, A.Y. (2012). AnimalTFDB: a comprehensive animal transcription factor database. Nucleic Acids Res 40, D144–149.

Zhou, Q., Liu, M., Xia, X., Gong, T., Feng, J., Liu, W., Liu, Y., Zhen, B., Wang, Y., Ding, C., and Qin, J. (2017). A mouse tissue transcription factor atlas. Nat Commun 8, 15089.

